# Mutations Affecting Polar Nuclei Number, Antipodal Cell Cluster Size, Cellularization, and Nuclear Localization in Maize Embryo Sacs

**DOI:** 10.1101/2022.11.10.516006

**Authors:** Antony M. Chettoor, Bing Yang, Matthew M. S. Evans

## Abstract

The maize female gametophyte is comprised of four cell types: two synergids, an egg cell, a central cell, and a variable number of antipodal cells. In maize, these cells are produced after three rounds of free-nuclear divisions followed by cellularization, differentiation, and proliferation of the antipodal cells. Cellularization of the eight-nucleate syncytium produces seven cells with two polar nuclei in the central cell. Nuclear localization is tightly controlled in the embryo sac as evidenced by the regular, stereotypical position of the nuclei in all syncytial stages of female gametophyte development. This leads to precise allocation of the nuclei into the cells upon cellularization. Nuclear positioning within the syncytium is highly correlated with their identity after cellularization. Two mutants are described with extra polar nuclei, abnormal antipodal cell morphology, and reduced antipodal cell number, which is correlated with a frequent loss of auxin signaling in the antipodal cell cluster. Mutations in one of these genes, *indeterminate gametophyte2* encoding a MICROTUBULE ASSOCIATED PROTEIN65-3 homolog, shows a requirement for MAP65-3 in cellularization of the syncytial embryo sac and that the identity of the nuclei in the syncytial female gametophyte can be changed very late before cellularization.

## INTRODUCTION

The plant life cycle has genetically active diploid and haploid phases, called the sporophyte and gametophyte (Walbot and Evans, 2003). The Polygonum type of female gametophyte development is the most common in angiosperms. The female gametophyte is derived from a single meiotic product, the chalazal megaspore (or functional megaspore (FM)). After meiosis, the FM undergoes three rounds of free nuclear divisions to produce an eight-nucleate syncytium. After the first division the two nuclei migrate to opposite poles of the embryo sac and are separated by a central vacuole. The nuclei then undergo two more rounds of synchronous divisions to produce an 8-nucleate syncytium with micropylar and chalazal clusters of nuclei. The embryo sac then cellularizes to produce seven cells of four cell types: the two synergids, the egg cell, the central cell, and three antipodal cells (Evans and Grossniklaus, 2009; Skinner and Sundaresan, 2018; Erbasol Serbes et al., 2019; Hater et al., 2020). In maize the antipodal cells proliferate to produce a cluster of 20 to 100 cells, in contrast to those of Arabidopsis which do not divide.

One nucleus from each cluster migrates to the center of the female gametophyte to become the polar nuclei of the central cell. The migration and position of gametophyte nuclei are highly regular. Nuclear movement in the female gametophyte occurs in two phases: (1) movement of syncytial nuclei prior to cellularization, particularly the migration of the two nuclei to opposite poles after the first division, where they divide twice to form two clusters of four nuclei; and (2) movement of the two polar nuclei. Polar nuclei movement itself can be divided into two phases: (1) one nucleus from each of the micropylar and chalazal poles migrates to the midline of the central cell where they partially fuse in maize; and (2) the partially fused polar nuclei move to the micropylar end of the central cell next to the egg apparatus.

Many mutants have now been identified that disrupt progression through the syncytial phase of female gametophyte development (reviewed in (Erbasol Serbes et al., 2019)). In Arabidopsis, mutations in the microtubule nucleation component, GCP2, and double mutants in gamma-tubulin genes cause defects in both the male and female gametophyte development, most commonly causing disruption after the second free-nuclear division in the female gametophyte (Pastuglia et al., 2006; Nakamura and Hashimoto, 2009).

A few genes have also been identified that are critical for nuclear movement within the female gametophyte. Kinesin encoding genes, *AtNACK1/HINKEL* and *STUD/TETRASPORE/AtNACK2*, are also necessary for proper cellularization and nuclear positioning with frequent failure in cytokinesis and mislocalization of nuclei occurring as early as immediately before cytokinesis with eventual clustering of nuclei in the center of the embryo sac (Tanaka et al., 2004). Late phase movement of the polar nuclei can be disrupted by disruption of F-actin dynamics (Kawashima and Berger, 2015). Fusion of the two polar nuclei in Arabidopsis requires a HSP70-like gene IMMUNOGLOBULIN BINDING PROTEIN (BiP) genes, demonstrating a conserved nuclear fusion mechanism between plants and yeast (Maruyama et al., 2010). Several mutants in Arabidopsis also suggest a role for mitochondria in polar nuclei fusion, *gametophyte factor2, syco-1, gamete cell defective1*, and *nuclear fusion defective1* (Christensen et al., 2002; Kagi et al., 2010; Wu et al., 2012).

In maize, during the free nuclear phase, microtubules form radiate, perinuclear arrays with connections between the sister nuclei at the micropylar and chalazal poles of the 4- and 8-nucleate stages in addition to cortical arrays (Huang and Sheridan, 1994, 1996). Phragmoplasts do not form between nuclei during the free nuclear divisions, but, after the last free nuclear division, phragmoplasts form simultaneously between sister nuclei before cytokinesis. The role of the cytoskeleton in this process has previously been suggested based on its organization during megagametogenesis (Webb and Gunning, 1994). In supportof this model, Arabidopsis gamma-tubulin mutants have defects in cytokinesis and nuclear positioning (Pastuglia et al., 2006; Nakamura and Hashimoto, 2009).

The embryo sac is polarized along the micropylar-chalazal (M-C) axis with the egg cell and synergids at the micropylar end and the antipodal cells at the chalazal end of the embryo sac). Mutant analysis suggests that positional information in the embryo sac is important for cell identity, as if there are determinants with unequal distribution within the embryo sac (Evans, 2007; Pagnussat et al., 2007). In Arabidopsis, asymmetric distribution of the ER-localized CKI1 protein to the chalazal end of the syncytial female gametophyte and its subsequent incorporation into the central cell and antipodal cells are crucial for distinguishing central cell and egg cell fate (Yuan et al., 2016). In *cki1* mutants, central cells and antipodal cells express egg cell specific markers. Asymmetric distribution of auxin signaling within the nucellus also seems to be important for egg cell identity based on a variety of experiments either increasing or reducing auxin levels or signaling (Pagnussat et al., 2009; Ceccato et al., 2013; Lituiev et al., 2013; Panoli et al., 2015). Whether auxin signaling acts cell-autonomously in the female gametophyte or acts in the surrounding nucellus to regulate the female gametophyte indirectly through an additional signal is unclear.

In maize, the egg-cell expressed ZMEAL1 protein is critical for suppressing central cell fate in antipodal cells (Krohn et al., 2012). Several genes have been identified that are important in restricting gamete cell fate in the Arabidopsis embryo sac, including genes encoding spliceosomal components (Gross-Hardt et al., 2007; Moll et al., 2008) and homologs of centromere proteins (Kirioukhova et al., 2011). *AMP1* in Arabidopsis is required to prevent the acquisition of egg cell fate in synergids (Kong et al., 2015). Many other advances have been made in the determination of embryo sac cell identities. The *myb64* and *myb119* are important for restricting chalazal identity, and *myb119* is downregulated in *cki* mutants (Rabiger and Drews, 2013). *AGL80* and *AGL61* are critical for central cell specification and the expression of central cell specific genes like *DEMETER* (Portereiko et al., 2006; Bemer et al., 2008; Steffen et al., 2008).

RKD1 and 2 are critical for egg cell identity in Arabidopsis, and several RKD genes are preferentially expressed in egg cell and can promote egg cell identity (Koszegi et al., 2011; Chardin et al., 2014; Tedeschi et al., 2017). The role for RKDs in gamete development (both egg and sperm) was also seen in marchantia (Koi et al., 2016; Rövekamp et al., 2016). Egg cell fate is expanded in a number of mutants, loss-of-function of *wyrd, lachesis, clotho*, and *atropos* and ectopic expression of *BEL1-LIKE HOMEODOMAIN1* (Gross-Hardt et al., 2007; Pagnussat et al., 2007; Moll et al., 2008; Kirioukhova et al., 2011; Kong et al., 2015). In *verdandi* mutants synergids express antipodal cell markers and vice versa, suggesting a common mechanism for regulating the two poles of the embryo sac. Expression of synergid genes and filiform apparatus development require the *MYB98* gene (Kasahara et al., 2005; Punwani et al., 2007).

In maize, it had been proposed that the position of the nuclei is critical for their eventual fate after cellularization, based in part on the identity of extra cells formed in the *indeterminate gametophyte1* mutant (Lin, 1978, 1981; Guo et al., 2004). The formation of extra egg cells is often associated with abnormal positioning of syncytial female gametophyte nuclei in several Arabidopsis mutants, (Gross-Hardt et al., 2007; Pagnussat et al., 2007; Moll et al., 2008; Kirioukhova et al., 2011; Kong et al., 2015). The correlation of the fate of a nucleus in the syncytial embryo sac with its position has been demonstrated in the Arabidopsis egg apparatus (Sun et al., 2021). Repositioning one of the nuclei within the micropylar cluster by disrupting actin filaments, so that two are in the future egg zone, causes the expression of egg cell markers in both of the nuclei rather than one.

The fluorescent reporters for auxin signaling in maize: *pDR5::RFP*, as a transcriptional reporter of auxin levels, and *pPIN1::PIN1-YFP* fluorescent protein fusion expressed from its native promoter are expressed strongly and specifically in the antipodal cell cluster within the embryo sac (Lituiev et al., 2013; Chettoor and Evans, 2015). Both are also expressed in the integuments and the micropylar nucellus of ovules, but are not expressed within immature embryo sacs. Expression of *pDR5::RFP* and *pPIN1::PIN1-YFP* in antipodal cells is associated with antipodal cell proliferation (Chettoor and Evans, 2015). Within the ovule, maize *pHistoneH1B::HISTONE1B-YFP* is expressed in the sporophytic cells of the ovule (*e*.*g*. nucellus, integuments) but within the embryo sac itself is expressed in the antipodal cells but not in other embryo sac cells (Chettoor and Evans, 2015).

The maize mutant *indeterminate gametophyte1* (*ig1*) disrupts normal embryo sac development in several ways (Kermicle, 1971; Lin, 1978, 1981; Huang and Sheridan, 1996; Huang and Sheridan, 1998; Evans, 2007). In *ig1* mutant embryo sacs, the early defects include excess rounds of free-nuclear divisions prior to cellularization producing embryo sacs with more than 8 nuclei (many of these have simple extra rounds of doubling but there can also be asynchronous nuclear divisions) (Lin, 1978, 1981; Huang and Sheridan, 1996). The position of these nuclei is also irregular, and, when cellularization occurs around these nuclei, the cells differentiate according to their positions. Consequently, mature *ig1* mutant embryo sacs contain extra egg cells, extra central cells, and extra polar nuclei within central cells. Because of their abnormal structure, many of these defective embryo sacs give rise to seeds with a variety of abnormalities including polyembryony, heterofertilization, miniature endosperms, early abortion of seeds, and the production of uniparental haploids (Kermicle, 1969, 1971). The later defects in morphology have been interpreted to be an indirect result of the earlier defects in proliferation.The extra polar nuclei appear to function normally as demonstrated by their effects on parental dosage in the endosperm, causing a deviation from the normal 2:1 ratio of maternal to paternal genomes. The miniature and aborted endosperm phenotypes arise from embryo sacs with extra polar nuclei (3 giving rise to miniature and 4 or more giving rise to aborted endosperms) (Lin, 1984). While pollination of wild-type diploid females by tetraploid males uniformly produces aborted kernels, pollination of diploid *ig1* mutant females by tetraploid males produces a few normal kernels with hexaploid endosperms with a 4 maternal:2 paternal genome ratio. Here we describe two mutants, *indeterminate gametophyte2* (*ig2*) and *ig3*, that also produce extra polar nuclei but through defects in embryo sac cellularization rather than through an excess of free nuclear divisions.

The *ig2* gene encodes a MICROTUBULE ASSOCIATED PROTEIN65-3/PLEIADE (MAP65-3/PLE) homolog. In Arabidopsis, MAP65-3 is required for cytokinesis and is localized to the phragmoplast (Muller et al., 2004; Ho et al., 2011; Sasabe et al., 2011). The MAP65-3 protein binds to antiparallel microtubules, stabilizing the mitotic spindle and promoting cytokinesis, possibly by allowing delivery of vesicles to the future cell plate. No role for *MAP65-3/PLE* in embryo sac development has been reported, and the *ple* mutations have no effect on fertility (Muller et al., 2002). In Arabidopsis MAP65-3 protein is present at other stages of the cell cycle but no function is yet ascribed to its presence there. Our data demonstrates a requirement for MAP65-3 (and hence cross-linking of anti-parallel microtubules) for cellularization of the syncytial embryo sac (and possibly the syncytial endosperm).

## RESULTS

### Identification of mutants

Active *Mutator* transposon lines were screened for heterozygous mutants with phenotypes similar to or overlapping with *ig1*. In particular ears were screened for reduced fertility and the production of miniature and aborted kernels despite being pollinated by homozygous wild-type plants. From this screen, two mutants were identified, named *indeterminate gametophyte2-O* (*ig2-O*) and *ig3-O* from the initial resemblance to *ig1*.

Pollination of either of these mutants as heterozygous females with wild-type pollen produced ears with partial sterility, as well as a number of seed defects – primarily aborted kernels and viable seeds with miniature endosperms (Figure 1 and Supplementary Figure 1, and Table 1 and 2). The most distinctive phenotype is the production of viable, miniature kernels that are hardly ever seen in wild-type females. There is no effect on seed phenotypes from male transmission of either *ig2-O* or *ig3-O* with similar percentages of rare abnormal kernel types as wild type, confirming the maternal basis of the effects of these mutations on seed development.

**Table 1.** Seed phenotypes caused by *ig2-O*.

| Table 1. Seed phenotypes caused by *ig2-O* |  |  |  |  |  |  |  |  |
| --- | --- | --- | --- | --- | --- | --- | --- | --- |
|  | Normal endosperm |  |  | Miniature endosperm |  |  | Aborted endosperm/<br>empty pericarp | Etched, or pitted endosperm |
| Inbred background and type of cross | Normal embryo | Abnormal or aborted embryo | Twins | Normal embryo | Abnormal or absent embryo | Twins |  |  |
| Wild type female by wild type male |  |  |  |  |  |  |  |  |
| +/+ x +/+ (1346 seeds) | 98.3% | 0.2% | 0 | 0 | 0 | 0 | 1.0% | 0.5% |
| Wild type female by *ig2* heterozygous male |  |  |  |  |  |  |  |  |
| +/+ x +/*ig2-O* (419 seeds) | 97.6% | 0.5% | 0 | 0 | 0 | 0 | 1.2% | 0.7% |
| *ig2* heterozygous female by wild type male |  |  |  |  |  |  |  |  |
| +/*ig2-O*; A158 x +/+ (346) | 93.6% | 0 | 0 | 2.3% | 0 | 0 | 4.1% | 0 |
| +/*ig2-O*; B73 x +/+ (1453) | 88.5% | 0.8% | 0.1% | 5.4% | 0.2% | 0 | 5.0% | 0 |
| +/*ig2-O*; Mo17 x +/+; (301) | 96.0% | 0 | 0 | 1.7% | 0 | 0 | 1.3% | 1.0% |
| +/*ig2-O*; W23 x +/+ (1247) | 97.0% | 0.5% | 0 | 1.9% | 0 | 0 | 0.6% | 0 |
| *ig2* heterozygote self-pollination |  |  |  |  |  |  |  |  |
| +/*ig2-O*; Mo17 x +/*ig2-O* (389) | 75.1% | 0.3% | 0 | 0.5% | 0 | 0 | 24.2% | 0 |

**Table 2.** Seed phenotypes caused by *ig3-O*.

| **Table 2. Seed phenotypes caused by *ig3-O*** |  |  |  |  |  |  |  |  |
| --- | --- | --- | --- | --- | --- | --- | --- | --- |
|  | Normal endosperm |  |  | Miniature endosperm |  |  | Aborted endosperm/<br>empty pericarp | Etched, or pitted endosperm |
| Inbred background and type of cross | Normal embryo | Abnormal or aborted embryo | Twins | Normal embryo | Abnormal or absent embryo | Twins |  |  |
| Wild type female by *ig3* heterozygous male |  |  |  |  |  |  |  |  |
| *+/+* x *+/ig3-O* (1847 seeds) | 97.9% | 0 | 0 | 0.1% | 0.1% | 0 | 1.1% | 0.8% |
| *ig3* heterozygous female by wild type male |  |  |  |  |  |  |  |  |
| *+/ig3-O*; B73 x *+/+* (2885) | 86.3% | 4.4% | 0.1% | 3.4% | 1.0% | 0 | 4.7% | 0.2% |
| *+/ig3-O*; Mo17 x *+/+* (511) | 93.7% | 1.3% | 0 | 2.4% | 0.6% | 0 | 2.0% | 0 |
| *+/ig3-O*; W23 x *+/+* (1769) | 92.4% | 0.8% | 0.1% | 5.3% | 0.8% | 0 | 0.6% | 0 |
| *ig3* heterozygote self-pollination |  |  |  |  |  |  |  |  |
| *+/ig3-O*; Mo17 x *+/ig3-O* (786) | 83.7% | 4.7% | 0 | 1.7% | 2.1% | 0 | 7.8% | 0 |
| *+/ig3-O*; W23 x *+/ig3-O* (580) | 75.6% | 6.9% | 0 | 8.3% | 7.9% | 0 | 4.1% | 0.2% |
| *ig3* homozygote self-pollination |  |  |  |  |  |  |  |  |
| *ig3/ig3*; A158 x *ig3/ig3* (304) | 50.0% | 2.6% | 0 | 9.2% | 9.2% | 0 | 29.0% | 0 |

**Figure 1.**
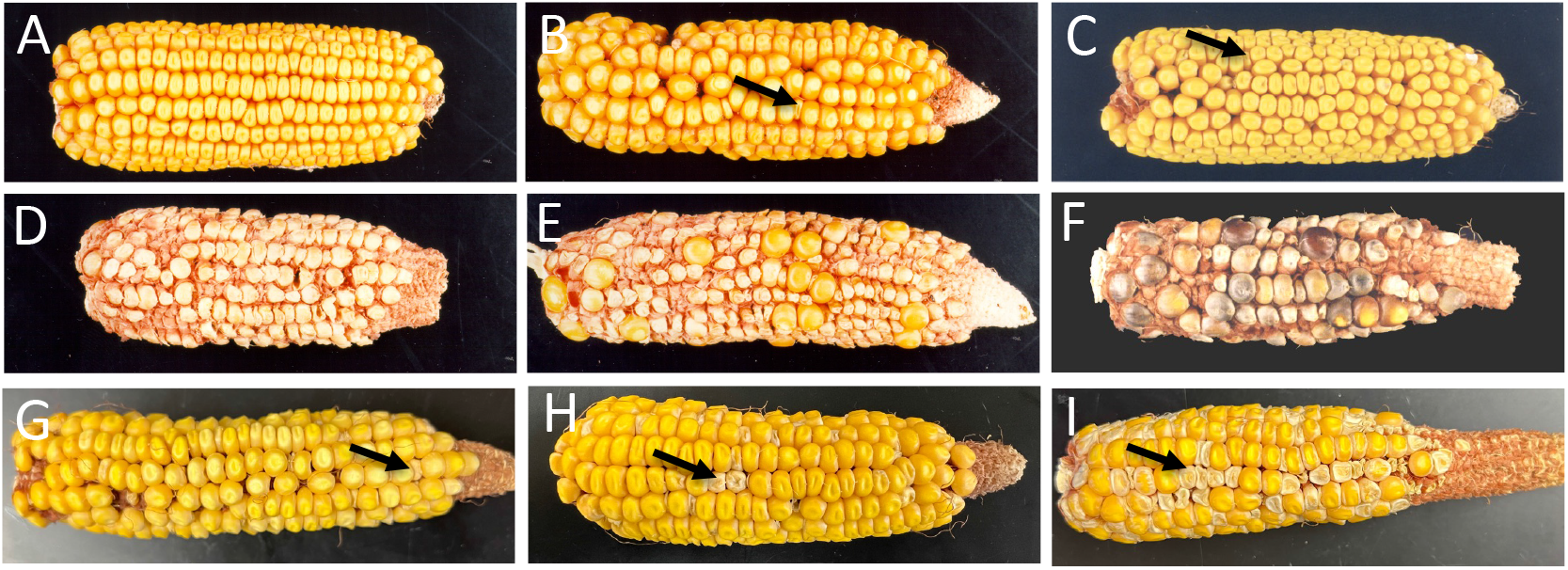
Seed phenotypes of *ig2-O* and *ig3-O* females. (A) wild-type self-pollination. (B) *ig2-O/+* female x wild-type pollen, (C) *ig3-O*/*+* female x wild-type pollen (D) Wild type female pollinated by tetraploid male. (E) *ig2-O/+* pollinated by tetraploid male. (F) *ig3-O/+* pollinated by tetraploid male. (G) *ig2-O/+* self-pollination. (H) *ig3-O/+* self-pollination. (I) *ig3-O/ig3-O* self-pollination. arrows=miniature kernels.

The *ig2* gene was initially rough mapped to the long arm of chromosome 8 between the Simple Sequence Repeat (SSR) marker *bnlg666* and the indel marker *IDP6826*. This map location was confirmed by linkage to the defective kernel mutant, *floury3* (Nelson and Ma, 1975; Li et al., 2017). The *ig3* gene was rough mapped to chromosome 2 between the SSR markers *bnlg1448* and *bnlg1036*, likely on the short arm of chromosome 2 near the centromere.

### Effects on seed development and mutant allele transmission

In wild type, pollination of wild-type diploid females by tetraploid males causes all of the kernels to abort development early because of the 2 maternal : 2 paternal (2m : 2p) genome ratio in the endosperm (deviating from the normal 2 maternal : 1 paternal ratio) (Figure 1B). In *ig1*, the presence of extra polar nuclei in the central cells causes some of the kernels from diploid *ig1* females pollinated with tetraploid males to develop normally, because the embryo sacs have four polar nuclei leading to a 4 maternal : 2 paternal genome ratio in the endosperm thus restoring the normal 2 : 1 ratio (Lin, 1978, 1981, 1984). To test if the same phenomenon may be occurring in *ig2* and *ig3*, mutant heterozygous diploid females were crossed by wild-type tetraploid males. In both *ig2* and *ig3* mutants, some fully developed viable kernels were produced in this cross (Figure 1D,E and Supplementary Figure 1). This is consistent with a subset of the embryo sacs having tetraploid central cells, possibly by containing four polar nuclei like *ig1* mutants rather than the two polar nuclei of wild type. These results are consistent with the fertilization of mutant embryo sacs with 3 polar nuclei and 4 polar nuclei producing the miniature kernels and aborted kernels, respectively, by haploid pollen. *ig1* mutants produce miniature and aborted kernels for the same reason (Lin, 1984).

The crosses of *ig2-O/+* and *ig3-O/+* mutants by tetraploid males also produced some kernels with endosperm development intermediate between the miniature and empty pericarp phenotypes not seen in crosses of wild type by tetraploid males. One explanation is that these result from fertilization of triploid central cells by diploid pollen grains (3m : 2p genome ratios) which is closer to the normal 2 maternal : 1 paternal ratio in the endosperm than either the 2m : 2p or 4m : 1p that result in endosperm abortion. In crosses of *ig1* by tetraploid males, these 3m : 2p endosperms developed normally for more days than 2m : 2p endosperms before eventually deviating from kernels with a ratio of 2m : 1p (Lin, 1984).

The frequency of abnormal kernels varied with genetic background. The suppression of the mutant phenotype can be seen most clearly from the percentage of normal seeds from mutant heterozygous. The strongest phenotypes for *ig2-O* occurred in B73 backgrounds, while the strongest suppression was produced in Mo17 and W23 inbred lines. For *ig3-O*, the strongest phenotype was seen in B73, but unlike *ig2-O* the next strongest phenotypes were seen in W23 and Mo17 with suppression in W64A. The frequency of both miniature kernels and empty pericarp/aborted endosperm phenotypes reached as high as ∼5% in *ig2-O/+* heterozygotes crossed by wild type, with similar maximum frequencies (although in different inbred backgrounds) in *ig3-O*. In *ig3-O* there is a shift in the frequencies of miniature vs. aborted kernels depending on genetic background with a decrease in the empty pericarp class in *ig3-O* but an increase in the miniature kernel class in W23 relative to B73. One interesting distinction between the two mutants is that *ig3-O* more frequently affects embryo development than does *ig2-O*.

Examination of ears from mutant heterozygotes revealed a range of partial sterility depending upon genetic background (Table 3), reaching as high as 28% for *ig2-O/+* and 26% for *ig3-O/+*. This level of sterility indicates that up to half of the mutant embryo sacs fail to produce a seed. The transmission of the *ig2-O* and *ig3-O* mutant alleles through both male and female gametophytes was tested to determine if the maternal effect on seed development was indeed gametophytic and then to determine whether the mutations affect only the female or affect both male and female. The rate of *ig2-O* and *ig3-O* male transmission is not statistically different from the wild-type alleles indicating no detrimental effects of either on the pollen (Table 4).

**Table 3.** Reduced fertility of *ig2-O/+* and *ig3-O/+* Heterozygotes.

| **Table 3. Reduced fertility of *ig2-O/+* and *ig3-O/+* heterozygotes** |  |
| --- | --- |
| Inbred background and type of cross | Seedless ovules |
| *+/+* x *+/+* (1371 ovules) | 1.8% |
| *+/ig2*; B73 x *+/+* (794 ovules) | 28.1% |
| *+/ig2*; Mo17 x *+/+* (347 ovules) | 13.3% |
| *+/ig2*; W23 x *+/+* (1330 ovules) | 6.2% |
| *+/ig3*; B73 x *+/+* (2249 ovules) | 25.7% |
| *+/ig3*; Mo17 x *+/+* (576 ovules) | 24.5% |
| *+/ig3*; W23 x *+/+* (489 ovules) | 9.0% |

**Table 4.** Transmission of *ig2-O* and *ig3-O* mutations to normal sized seeds.

| Table 4. Transmission of *ig2-O* and *ig3-O* mutations to normal sized seeds |  |  |
| --- | --- | --- |
| Cross | *ig2-O/+* heterozygotes | Homozygous wild type |
| +/+ female x +/*ig2-O* male | 83* | 99* |
| +/*ig2-O* female x +/+ male | 175*† | 358*† |
|  | *ig3-O/+* heterozygotes | Homozygous wild type |
| +/+ female x +/*ig3-O* male | 42 | 46 |
| +/*ig3-O* female x +/+ male | 83† | 144† |
| *Some of these are based on transmission of the tightly linked *floury3* mutation in repulsion to *ig2-O* |  |  |
| †Significantly different from 50% at p<0.01 |  |  |

Both mutations, on the other hand, show reduced transmission through the female with approximately one third of the normal kernels inheriting the mutant allele (Table 4). This is important to show that the effects of the mutations on the production of abnormal kernels and the reduction of fertility are gametophytic, *i. e*. abnormal kernels and reduced fertility are caused by action of the mutation in the embryo sac rather than a dominant action of the mutation in the diploid tissues of the sporophyte, such as the nucellus or integuments. Interestingly, the actual deficit of the mutant allele in the progeny matches well with the predicted frequency based on the percentage of sterile ovules in the mutant heterozygotes (Table 3).

In addition to their roles in the embryo sac that lead to seed abnormalities and reduced fertility, both genes appear to have roles in the sporophyte as well. In self-pollinations of mutant heterozygotes, there is an increase in abnormal seed phenotypes (Tables 1 and 2). For *ig2*, there is a dramatic increase in the frequency of the aborted, empty pericarp kernel class even in a genetic background that suppresses the maternal effects of the mutation, where approximately ¼ of the seeds abort early. In rare cases there is enough endosperm development to foster partial progression of embryo development and seed germination. The mutant endosperms undergo more extensive cell death in the central starchy endosperm than is seen in wild type, and the embryos are much smaller and abnormal in shape (Figure 2). The presumed *ig2-O/ig2-O* seedlings are highly abnormal. They have disorganized cell files in the leaves and scutellum and have irregularly shaped, bulbous epidermal cells (Figure 2D,E). These abnormal seedlings do not progress beyond the seedling stage, and mature homozygotes have never been recovered.

**Figure 2.**
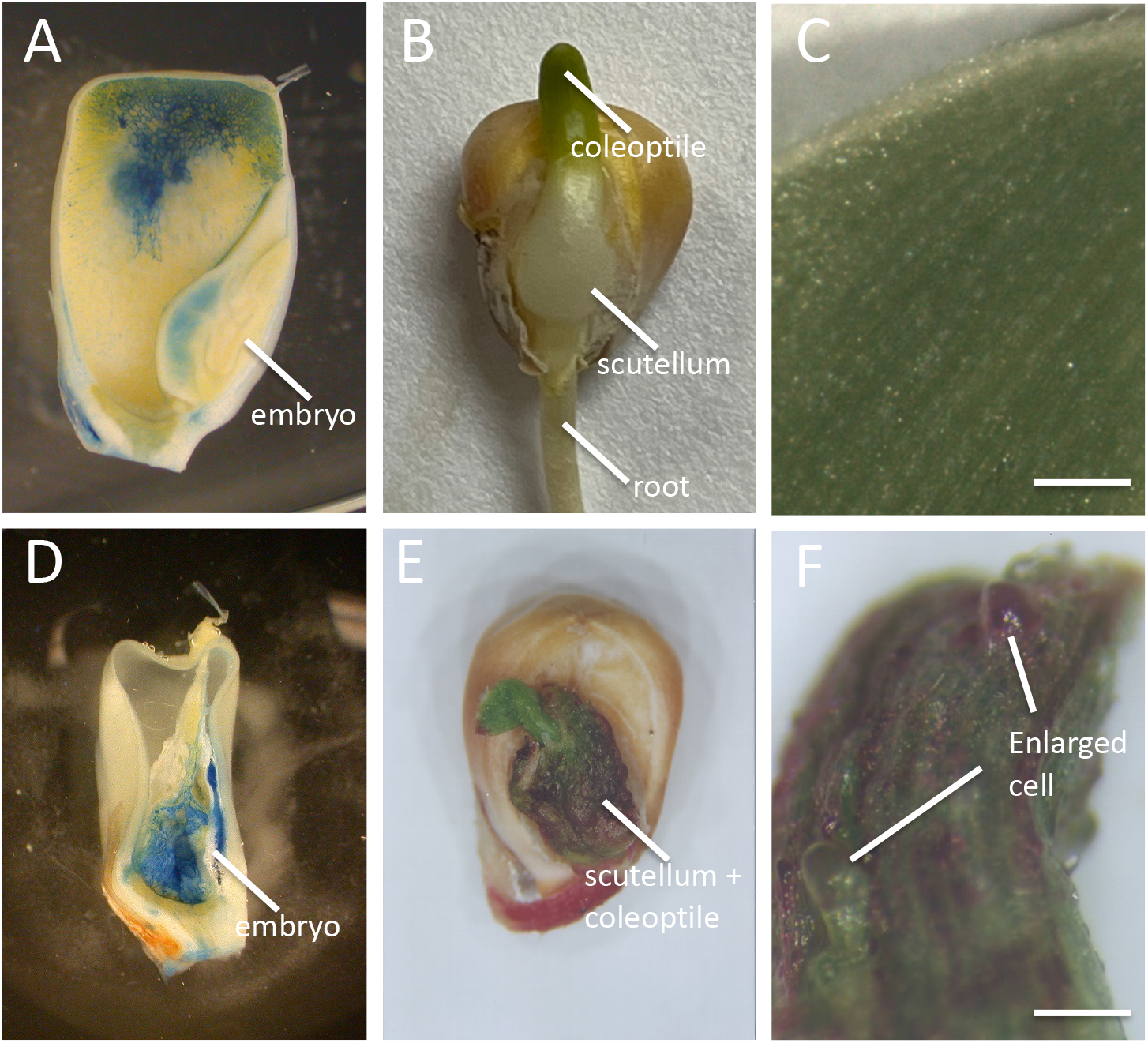
Phenotype of Homozygous *ig2-O/ig2-O* mutants. (A-C) Wild type. (D-F) *ig2-O/ig2-O*. (A, D) Sagittal section of 35 day after pollination kernel stained with Evans Blue to show cell death/programmed cell death in the Central Starchy Endosperm. Note that the entire *ig2* mutant endosperm has undergone cell death and the embryo is very small and underdeveloped. (B, D) Germinating seedling note the disorganized scutellum and coleoptile of the *ig2* mutant seedling. (C, F) Seedling leaf epidermis. The *ig2* mutant has less well defined cell files and enlarged bulbous cells on the epidermis that do not occur in wild type.

For *ig3-O*, there is a broader range of phenotypes for the presumed homozygotes. There is an increase in several different kernel classes in the self-pollinations, including the empty pericarp class and the miniature endosperm class, as well as an increase in the frequency of embryo defects (Table 2). In contrast to *ig2, ig3* homozygotes also can progress beyond seed development. The homozygous *ig3/ig3* seeds that germinate produce plants that are indistinguishable from wild type, except that they produce abnormal kernels at a higher frequency than *ig3*/*+* heterozygotes with only half of the seeds developing normally (Figure 1I and Table 2). Additionally, some of the small kernel types in *ig3-O* also have a loose pericarp, rather than a classic miniature phenotype (which is smaller than wild type but otherwise normal), although it is difficult to tell if genetic background may be contributing to some of this distinction (Figure 1H,I).

### Effects on embryo sac morphology

*ig2-O* causes a variety of defects in mature embryo sacs (Figure 3 and Table 5). These defects include all embryo sac cell types with changes in the morphology or number of: antipodal cells, central cells, egg cells and synergids (Table 5). Mature *ig2* embryo sacs can have extra polar nuclei (Fig. 3C) or extra central cells (Figure 3E), although these appear otherwise normal. Some of the most common effects are on the antipodal cells (Fig. 3 B,C,D) with over half of the mutant embryo sacs having antipodal cell cluster defects (Table 5). In wild-type embryo sacs, the antipodal cells are typically 20 to 100 cells in number and usually more densely cytoplasmical than other embryo sac cells depending on genetic background (Figure 3A and 3A) (Evans and Grossniklaus, 2009). Mutant embryo sacs often have fewer antipodal cells than wild type and these embryo sacs are frequently much larger and more vacuolated than those in wild type (Fig. 3B), although sometimes the antipodal cells are normal in appearance despite being fewer in number (Figure 3C). In some cases, there are no antipodal cells at all (Figure 3E). The number of cells in the egg apparatus – the synergids and egg cells – are also frequently reduced in *ig2* embryo sacs (Table 5). In the embryo sacs in which they form, the egg cells and synergids appear morphologically normal. In the most severe cases the *ig2* embryo sac consists of a single large cell with all of the nuclei clustered in the center (Fig. 3F).

**Table 5.** Summary of Mature Embryo Sac Phenotypes from *ig2-O/+* heterozygotes.

|  |  |  |  |  |  |  |  |  |  |  |  |  |  |  |  |  |
| --- | --- | --- | --- | --- | --- | --- | --- | --- | --- | --- | --- | --- | --- | --- | --- | --- |
| Normal (99 total) | Abnormal (80 total)* |  |  |  |  |  |  |  |  |  |  |  |  |  |  |  |
|  | Aborted/<br>Arrested | 1<br>Big<br>Cell | Antipodal Cell Phenotypes |  |  |  |  | Egg Apparatus Phenotypes |  |  |  |  | Central Cell Phenotypes |  |  |  |
|  |  |  | Fewer AC |  |  |  | wt<br>AC | Fewer Egg + Syn |  |  |  | Normal<br>Syn +<br>Egg | Normal<br>CC | Extra<br>PN | Extra<br>CC | 1<br>PN |
|  |  |  | Fewer<br>but<br>normal | No<br>AC | Small but<br>vacuolated | Large |  | 0<br>Syn<br>0<br>Egg | 0<br>Syn<br>1<br>Egg | 1<br>Syn<br>1<br>Egg | 1<br>Syn<br>0<br>Egg |  |  |  |  |  |
| 99 | 3 | 6 | 29 | 7 | 6 | 8 | 20 | 17 | 11 | 13 | 2 | 15 | 1 | 63 | 3 | 1<sup>†</sup> |
| *Some abnormal embryo sacs are in multiple categories and in some embryo sacs the whole embryo sac could not be visualized.<br><sup>†</sup>These may be wild-type embryo sacs in which the two polar nuclei have completed fusion during maturation, which occurs in a subset of wild-type embryo sacs depending on genetic background. |  |  |  |  |  |  |  |  |  |  |  |  |  |  |  |  |

**Figure 3.**
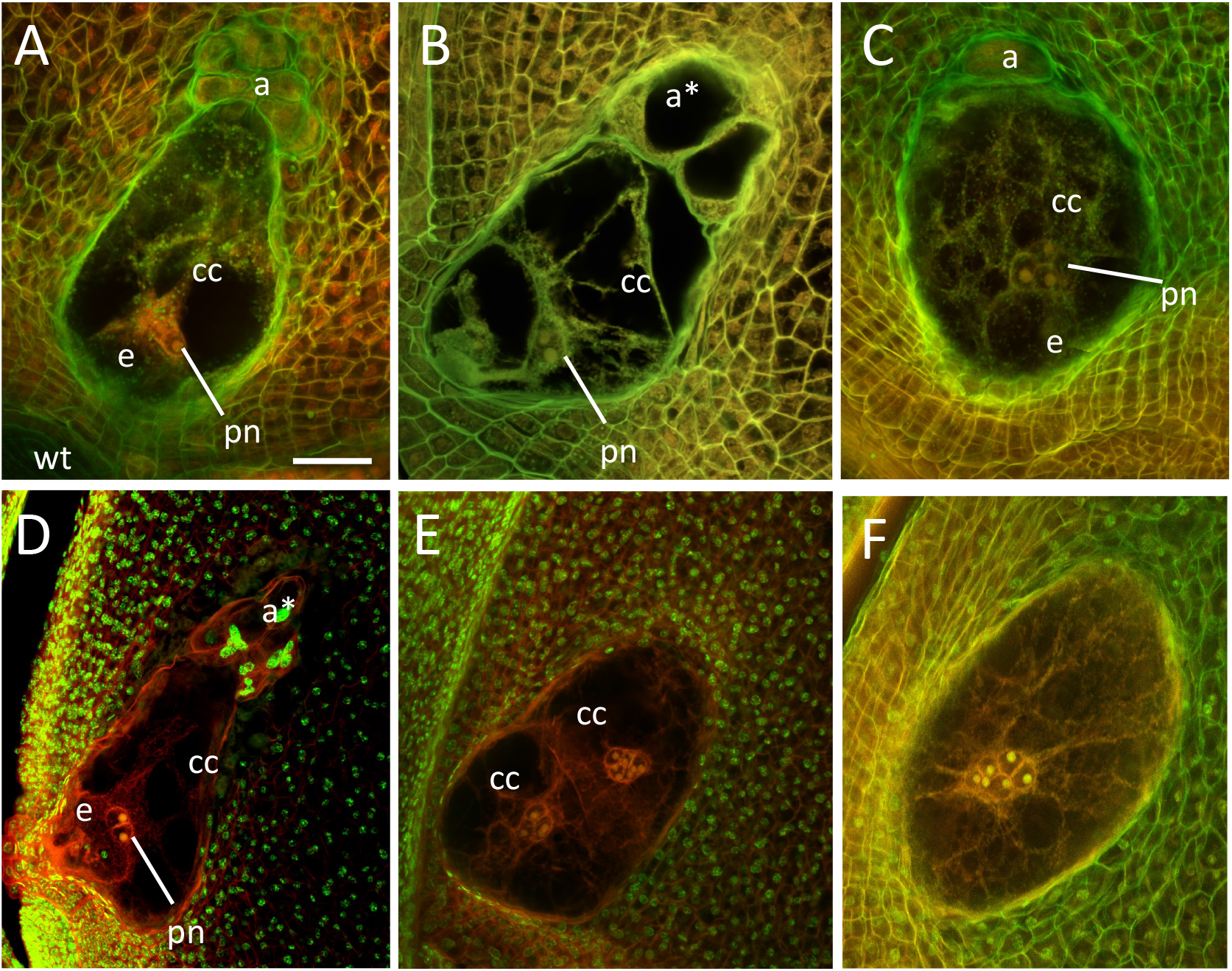
Effect of *ig2* on mature embryo sac morphology. (A) wild-type, (B-F) *ig2* embryo sacs. (B) *ig2* embryo sac with large vacuolated antipodal cells. (C) *ig2* embryo sac with one antipodal cell and excess polar nuclei. (D) *ig2* embryo sac with giant nuclei in the antipodal cells. (E) *ig2* embryo sac with multiple central cells each with extra polar nuclei (F) *ig2* embryo sac with a single large cell with many nuclei in the center. cc=central cell. e=egg cell. a=antipodal cells; pn=polar nuclei; a*=abnormally large, vacuolated antipodal cells. Scale bar = 50 μm.

Antipodal cell nuclei have a different morphology than the other embryo sac nuclei. In the syncytial female gametophyte and in the synergids, egg cell, and central cell the nuclei have a prominent nucleolus and stain poorly with Propidium Iodide. This morphology first appears in the Megaspore Mother Cell and persists in all female gametophyte cells except for the antipodal cells (Supplementary Figure 2). The antipodal cell nuclei transition from this morphology to a nucleus with speckled Propidium Iodide staining and without a prominent nucleolus that more closely resemble the surrounding nucellular cells. In *ig2* mutant embryo sacs, the nuclei at the chalazal end/in the antipodal cells occasionally fail to undergo this transition (4 abnormal : 21 normal) (Figure 4).

**Figure 4.**
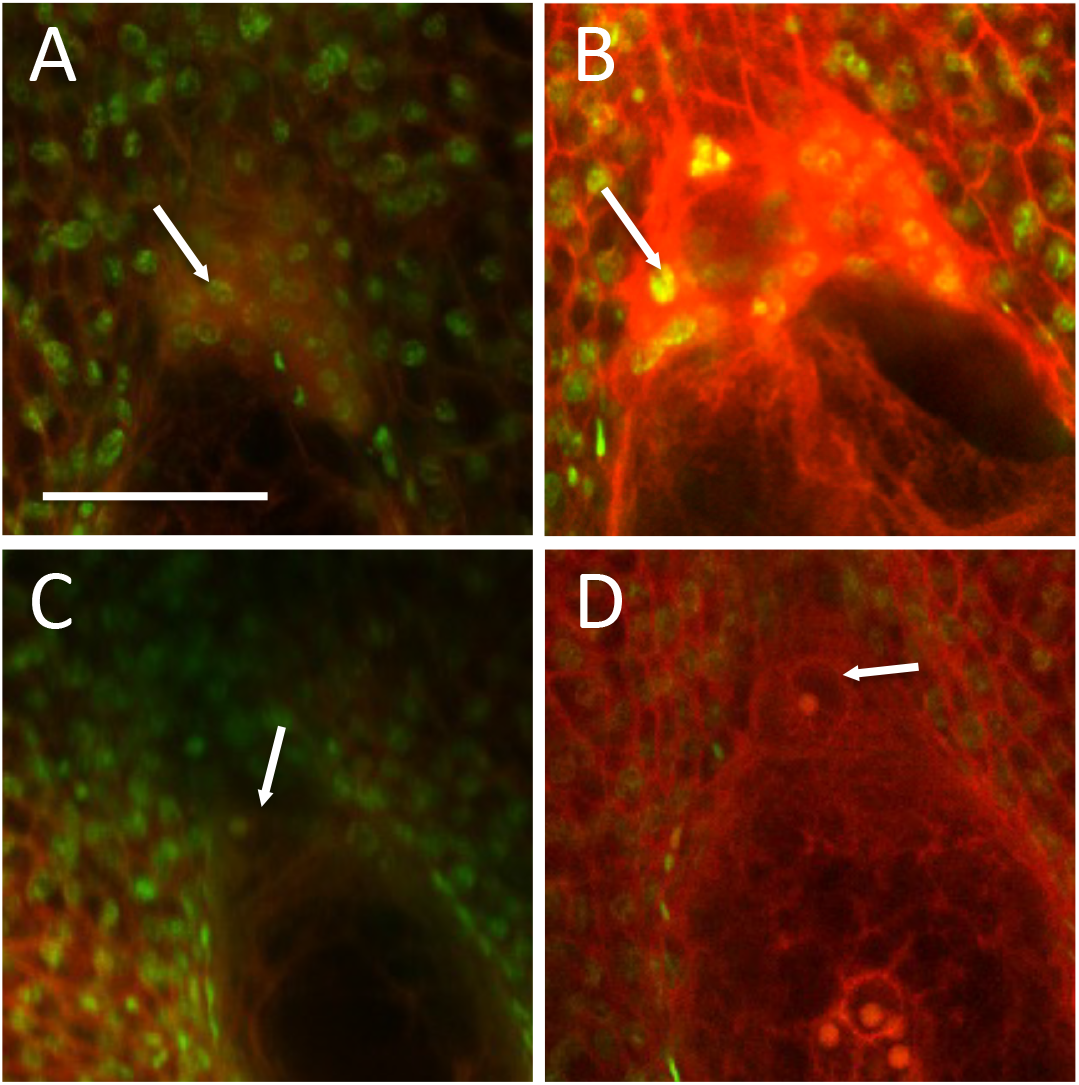
Effect of *ig2* on morphology of antipodal/chalazal nuclei. (A) wild-type notice the speckled staining with Propidium Iodide of the antipodal cell nuclei similar to the nucellular cells (B) *ig2* antipodal cells with normal nuclei. (C-D) *ig2* antipodal cell/chalazal nucleus with the prominent nucleolus and lack of Propidium Iodide staining normally seen in all embryo sac cells besides the antipodals. Scale bar = 50 μm.

Early stages of *ig2* embryo sac development appear indistinguishable from wild type (Supplementary Table 1). Normal embryo sac development includes three rounds of free nuclear division to produce an eight-nucleate syncytium with a cluster of 4 nuclei at each pole (Supplementary Figure 2). These embryo sacs were collected from immature maize ears which have a gradient of floral stages with the youngest at the tip of the ear and the oldest at the base of the ear. No abnormalities were detected in in 1-nucleate or 2-nucleate embryo sacs which have very uniform development. Some variability is seen in the nuclear positioning in 4-nucleate embryo sacs in both wild type and *ig2*. The two micropylar nuclei are almost always side by side oriented perpendicular to the micropylar-chalazal axis. The positioning of the chalazal nuclear pair is more variable but the frequency of the different configurations is the same in wild type and *ig2*. Half of the embryo sacs have the chalazal nuclei positioned where the line between them is perpendicular to the micropylar-chalazal axis. In the other half of the 4-nucleate embryo sac, these two nuclei are oriented parallel to the micropylar-chalazal axis. Additionally, in half of these embryo sacs the two nuclei are adjacent to one another (approximately as close together as the two chalazal nuclei are to each other when they are perpendicular to the long axis of the embryo sac). However, in the other half there is space between the two nuclei. The fractions of embryo sacs in these different categories are the same in wild type and *ig2*.

The first defect in *ig2* is seen at the transition between the 8-nucleate and early cellularized stages. In wild type, this transition occurs fairly quickly so that the 8-nucleate stage is less common than the other stages when looking at a developmental series of ovules (17 at 2-nucleate to 20 at 4-nucleate to 8 at 8-nucleate syncytial to 60 at an early cellularized stage) (Supplementary Table 1). In *ig2/+* heterozygotes, the 8-nucleate stage is seen more frequently suggesting the embryo sacs remain in this stage for more time in mutants (33 at 2-nucleate to 39 at 4-nucleate to 23 at 8-nucleate syncytial to 25 at an early cellularized stage). It is also around this stage that the first obvious defects appear. In wild-type, the two polar nuclei have migrated to the central region of the syncytium, usually along one of the side walls of the cell and may or may not be in contact already. In *ig2/+* heterozygotes, approximately 1/8^th^ of the embryo sacs (which should be 1/4^th^ of the mutant embryo sacs) have more than 2 nuclei in the central domain of the syncytial/early cellularized embryo sacs, suggesting that additional nuclei have failed to maintain their normal location at the syncytial poles at this stage (Figure 5). These nuclei are also commonly in the center of the cell rather than against one of the side walls. The developmental pattern of *ig2* mutant embryo sacs is summarized in Supplementary Figure 3.

**Figure 5.**
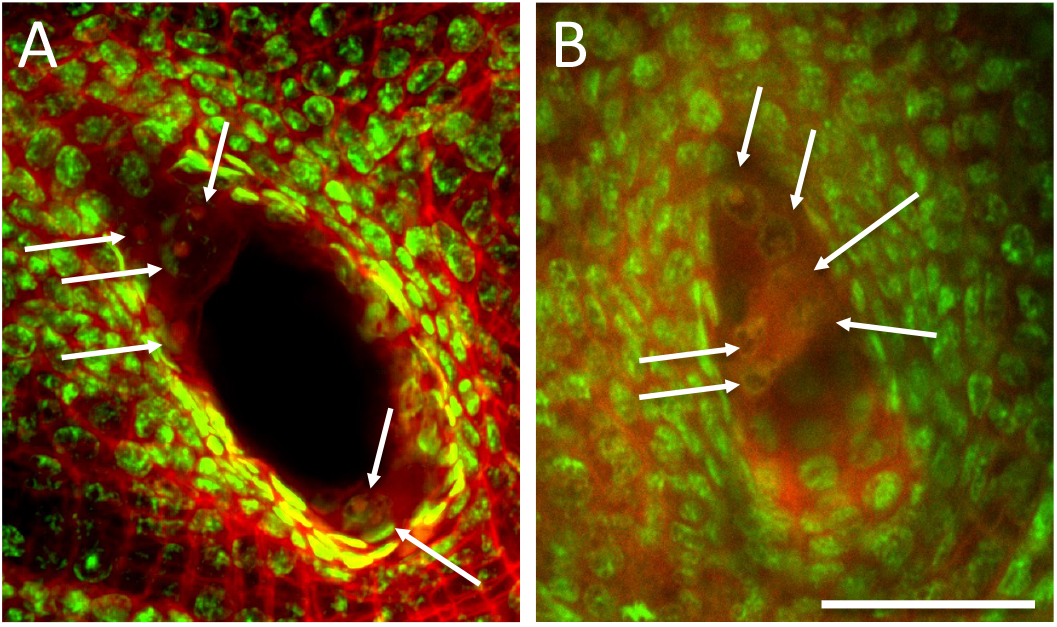
Effects of *ig2-O* on nuclear distribution at the time of cellularization. (A) Wild type. (B) *ig2-O* with nuclei in the center of the embryo sac. Arrows point to nuclei. Because not all of the nuclei are in the same plane in maize 8-nucleate embryo sacs only six nuclei are shown here. Scale bar = 50 μm.

Embryo sac defects caused by *ig3* overlap substantially with those caused by *ig2* (Table 6 and Figure 6). *ig3* mutant embryo sacs often have abnormal antipodal cell clusters. The antipodal cells can be fewer in number (Figure 6B,C,F) or absent (Figure 6E). Like in *ig2*, when *ig3* antipodal cells are fewer in number, they can have a normal morphology, be large and vacuolated, or small like wild type but less cytoplasmically dense. Like in *ig2*, some *ig3* mutant embryo sacs have extra polar nuclei (Fig. 6B). There are some phenotypes seen in *ig3* but not *ig2*. The polar nuclei in a few embryo sacs are located against one of the side walls of the central cell (Fig. 6F). Also, there are rare embryo sacs with abnormally shaped egg cells (Fig. 6B). The embryo sacs consisting of a single large cell seen in *ig2* have not been seen in *ig3*.

**Table 6.**
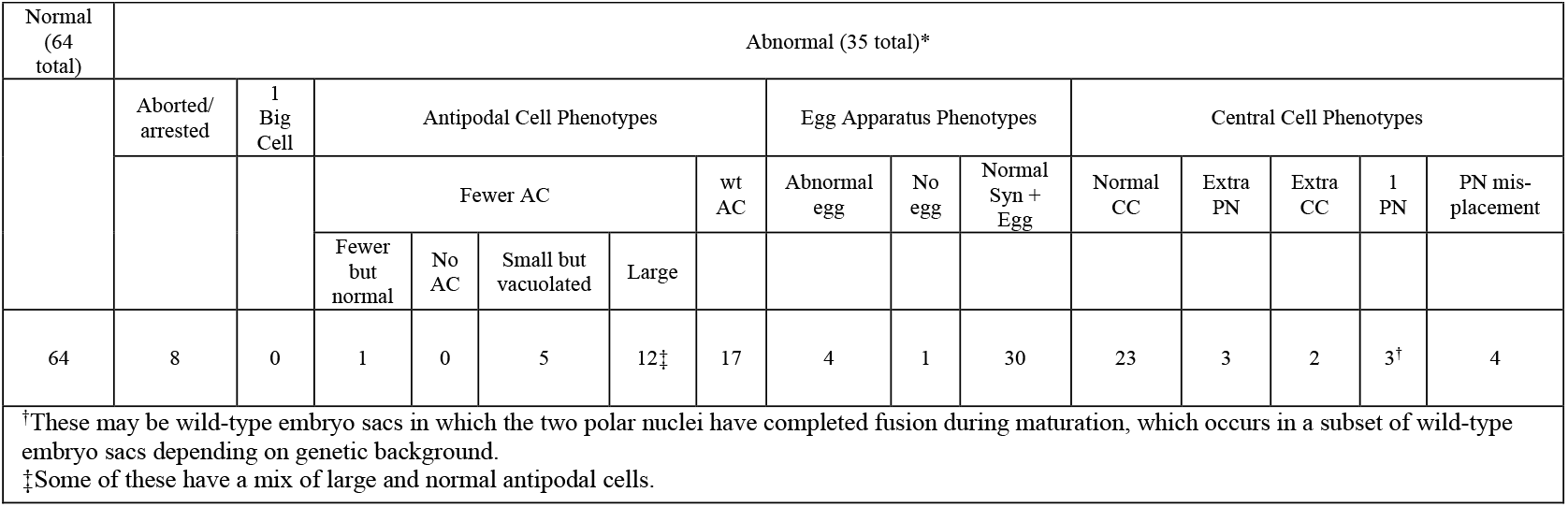
Summary of Mature Embryo Sac Phenotypes from *ig3-O/+* heterozygotes.

|  |  |  |  |  |  |  |  |  |  |  |  |  |  |  |  |
| --- | --- | --- | --- | --- | --- | --- | --- | --- | --- | --- | --- | --- | --- | --- | --- |
| Normal (64 total) | Abnormal (35 total)* |  |  |  |  |  |  |  |  |  |  |  |  |  |  |
|  | Aborted/<br>arrested | 1<br>Big<br>Cell | Antipodal Cell Phenotypes |  |  |  |  | Egg Apparatus Phenotypes |  |  | Central Cell Phenotypes |  |  |  |  |
|  |  |  | Fewer AC |  |  |  | wt<br>AC | Abnormal<br>egg | No<br>egg | Normal<br>Syn +<br>Egg | Normal<br>CC | Extra<br>PN | Extra<br>CC | 1<br>PN | PN mis-<br>placement |
|  |  |  | Fewer<br>but<br>normal | No<br>AC | Small but<br>vacuolated | Large |  |  |  |  |  |  |  |  |  |
| 64 | 8 | 0 | 1 | 0 | 5 | 12<sup>‡</sup> | 17 | 4 | 1 | 30 | 23 | 3 | 2 | 3<sup>†</sup> | 4 |
| <sup>†</sup>These may be wild-type embryo sacs in which the two polar nuclei have completed fusion during maturation, which occurs in a subset of wild-type embryo sacs depending on genetic background.<br><sup>‡</sup>Some of these have a mix of large and normal antipodal cells. |  |  |  |  |  |  |  |  |  |  |  |  |  |  |  |

**Figure 6.**
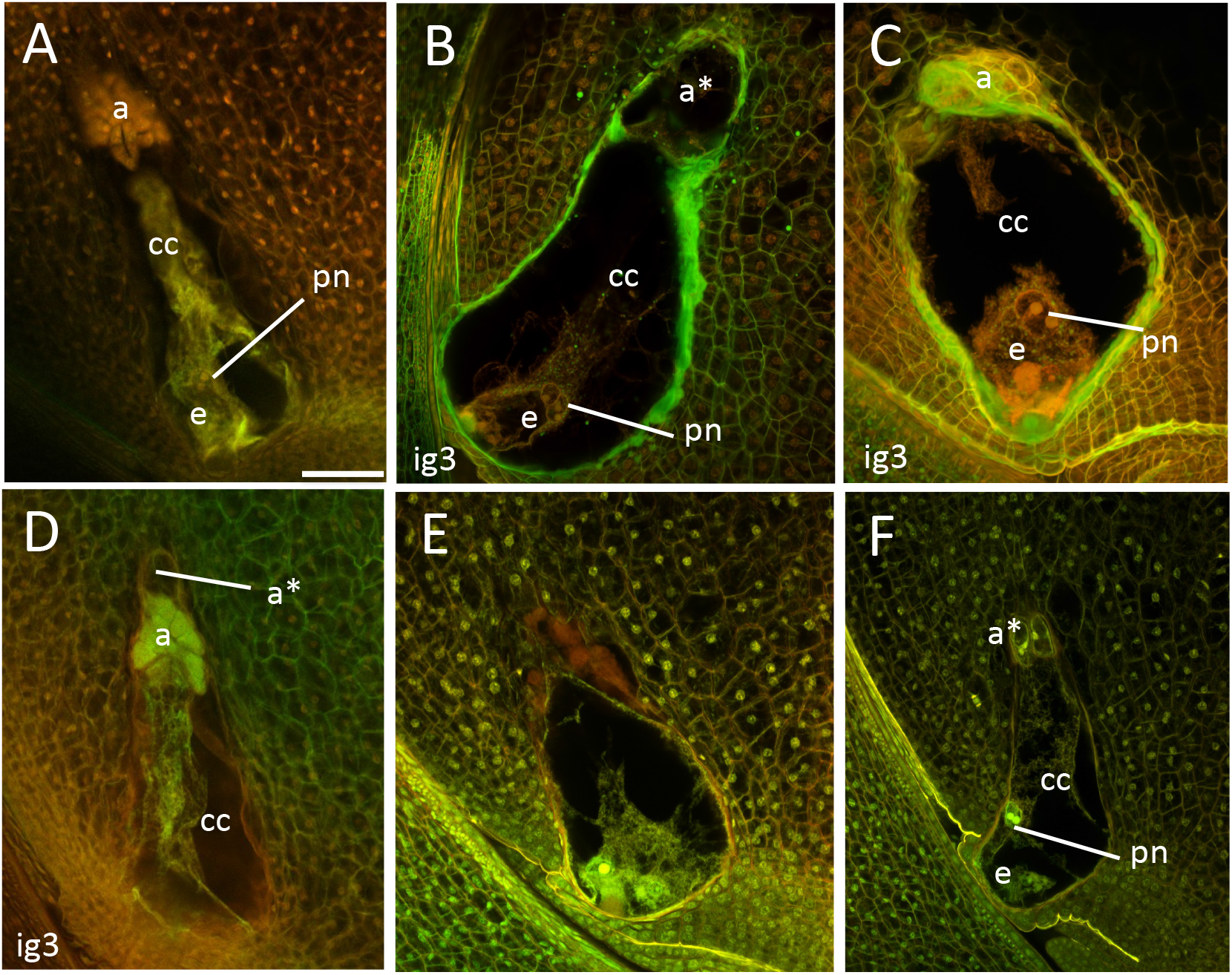
Effect of *ig3* on mature embryo sac morphology. (A) wild-type, (B-F) *ig3* embryo sacs. (B) *ig2* embryo sac with large vacuolated antipodal cells and three polar nuclei. (C) *ig3* embryo sac with one antipodal cell. (D) *ig3* embryo sac with a mix of normal and vacuolated antipodal cells. (E) *ig3* embryo sac with no antipodal cells (F) *ig3* embryo sac with misplaced polar nuclei against the side wall of the central cell. cc=central cell. e=egg cell. a=antipodal cells; pn=polar nuclei; a*=abnormally large, vacuolated antipodal cells. Scale bar = 50 μm.

Because of the frequent antipodal cell defects in *ig2* and *ig3*, fluorescent markers for the antipodal cells were crossed to both mutants to determine if any gene expression patterns normally associated with the antidopals are abnormal in the mutants. The fluorescent markers, *pHISTONE H1B(GRMZM2G164020)::HISTONE H1B-YFP, pDR5::RFP*, and *pPIN1(GRMZM2G098643*)*::PIN1-YFP*, are expressed in the antipodal cells but not in other embryo sac cells in maize (Lituiev et al., 2013; Chettoor and Evans, 2015). Expression of *pPIN1* and *pDR5* is absent in *Laxmidrib1* mutants with reduced antipodal cell numbers suggesting an association between antipodal cell proliferation and auxin signaling (Chettoor and Evans, 2015). Both *pPIN1* and *DR5* are only rarely expressed in the abnormal antipodal cells of *ig2* and *ig3* (Table 7 and Figure 7). Expression of *pHIS* is also usually absent from *ig2* mutant embryo sacs, but has not been tested in *ig3*. In no cases was there ectopic expression of these markers in other embryo sac cells. Since the mutant alleles and the transgenes are segregating in the embryo sacs and were crossed into the heterozygous plant from opposite parents, the possibility existed that the transgene (of unknown location) and the mutant allele could be linked and in repulsion. This would cause fewer of the mutant embryo sacs to inherit the transgene and so not express the transgene for that reason and not because of developmental defects caused by the mutation.

**Table 7.** Antipodal cell expression of fluorescent reporters in embryo sacs of plants hemizygous for the transgene and heterozygous for *ig2* or *ig3*.

|  |  |  |  |  |
| --- | --- | --- | --- | --- |
| Genotype of Plants | Normal ES without fluorescent reporter expression in antipodal cells | Normal ES with fluorescent reporter expression in antipodal cells | Abnormal ES without fluorescent reporter expression in antipodal cells | Abnormal ES with fluorescent reporter expression in antipodal cells |
| *ig2-O/+*;<br>*pPIN1::PIN1-YFP/-* | 40 | 44 | 25* | 6 |
| *ig2-O/+*;<br>*pDR5::RFP/-* | 23 | 20 | 24* | 4 |
| *ig2-O/+*;<br>*pHIS H1B::HIS H1B-YFP/-* | 54 | 54 | 23* | 8 |
| *ig3-O/+*;<br>*pPIN1::PIN1-YFP/-* | 17 | 22 | 7* | 0 |
| *ig3-O/+*;<br>*pDR5::RFP/-* | 39 | 36 | 16* | 0 |
| *Mutant embryo sacs included here are those with large antipodal cells which are easier to identify in the lower resolution live-cell imaging. Embryo sacs with normal antipodal cells or no antipodal cells were not included here. |  |  |  |  |

**Figure 7.**
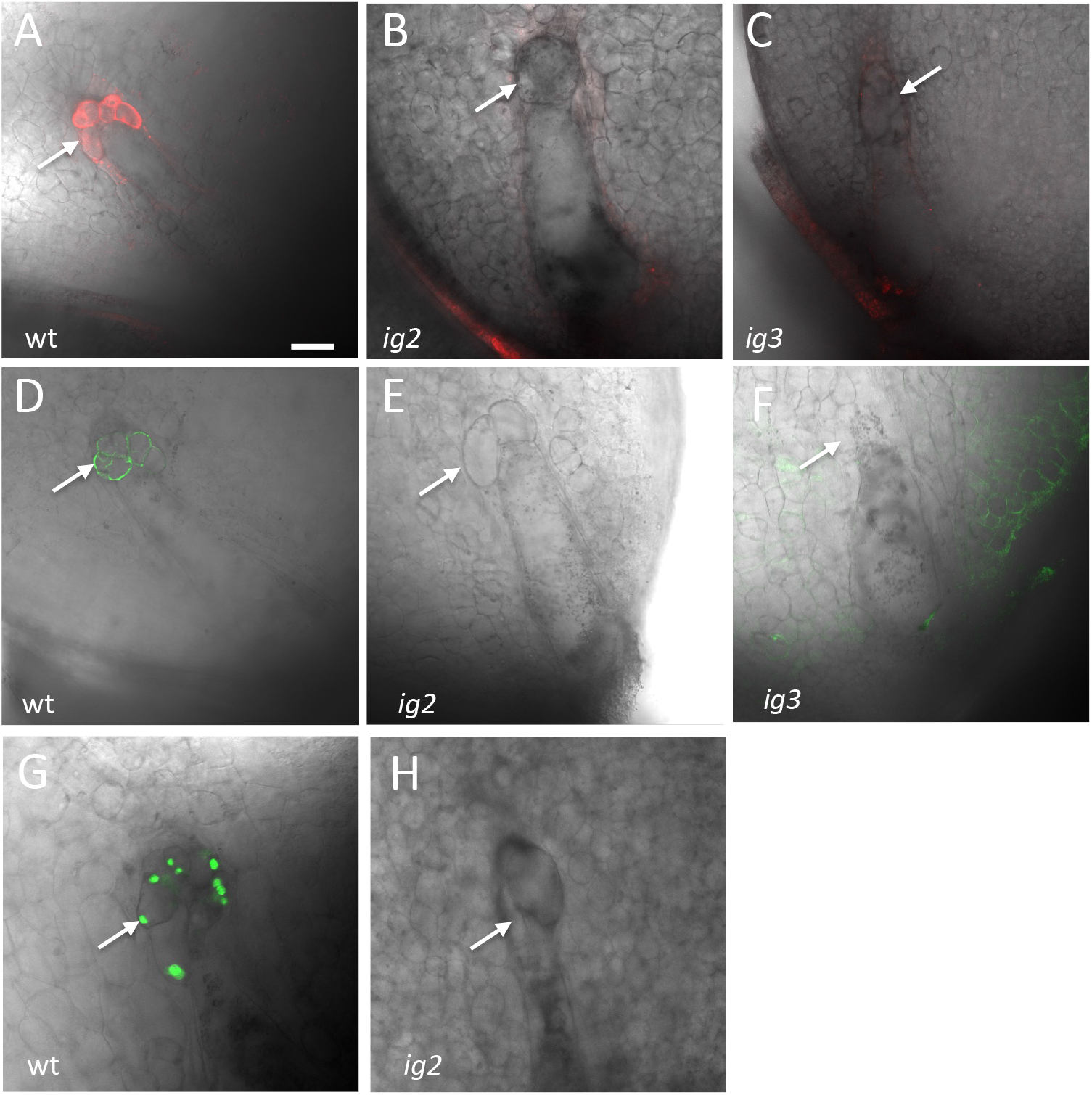
Effect of *ig2* on embryo sac expression of *pPIN1::PIN1-YFP* and*pDR5::RFP*. Live cell imaging of sibling (A, D, G) wild-type, (B, E, H) *ig2* embryo sacs with abnormal antipodal cells, and (C,F) *ig3* embryo sac with abnormal antipodal cells carrying either (A, B,C) *pDR5::RFP* or (D,E,F) *pPIN1::PIN1-YFP* or (G,H) *pHistoneH1B::HISTONEH1B-YFP*. Arrows indicate antipodal cell cluster. Scale bar = 50 μm.

However, linkage in repulsion would also cause the wild-type embryo sacs to be more likely to inherit the transgene and so more likely to express it. In all cases, about half of the wild-type embryo sacs express the transgene indicating that the mutant allele and the transgene are segregating independently and the reduced frequency of transgene expression is a consequence of the developmental abnormalities of the mutant embryo sacs and not an artifact of linkage (Table 7).

### Identification of the *ig2* gene

Since the *ig2* and *ig3* mutations arose in lines with active *Mutator* transposable elements, mutant lines were subjected to *Mu* Illumina sequencing to identify all of the *Mu* insertion sites in the genome for each of these mutations (Williams-Carrier et al., 2010). No *Mu* insertion sites were found in the *ig3* mutant lines that were not present in control non-mutant lines. However, the *Mu* Illumina sequencing revealed *Mu* insertions in two genes within the *ig2* containing interval on chromosome 8 in two separate *ig2* pedigrees; these genes are Zm00001d011615, encoding a MICROTUBULE ASSOCIATED PROTEIN65-3/PLEIADE (MAP65-3/PLE) homolog, and Zm00001d011878, encoding a BTB/POZ domain transcription factor. The *Mu* insertion in Zm00001d011615 is located in the 5’utr in the first exon. As is typical for *Mu* insertions (Barker et al., 1984), there is a target site duplication at the insertion site (Supplementary Figure 4). In Arabidopsis, *pleiade* mutant plants are viable but have defects in cytokinesis and produce enlarged cells in the root with multiple and/or enlarged nuclei (Muller et al., 2002; Muller et al., 2004). The latter phenotype is similar to the most severe phenotype seen in *ig2* embryo sacs consisting of a single large cell with multiple, centrally located nuclei.Consequently, to test if *ig2* is indeed this *MAP65-3* homolog, CRISPR-Cas9 was used to generate additional mutant alleles of Zm00001d011615.

A CRISPR-Cas9 construct was designed to target the coding region in the third exon of the Zm00001d011615 gene, and then transferred into the maize Hi-II genotype *via Agrobacterium* infection of immature embryos. The genome editing produced three independent, heritable mutant alleles. All were found in heterozygotes. No homozygous mutants were found, a fact which would be consistent with the predicted homozygous lethality of *ig2*. All three mutations contained a single base insertion at the same site but with different bases inserted, one each with an A, G, or T (Supplementary Figure 4). All three produced kernels with miniature or aborted endosperms at a similar frequency to *ig2-O*, allowing for effects of genetic background (Supplementary Table 2). Additionally, all three show non-complementation of the *ig2-O* homozygous lethality based on the increase in the frequency of the aborted kernel phenotype when heterozygotes for these alleles were crossed to *ig2-O*/+ heterozygous females (with similar frequencies as *ig2-O/+* self-pollinations). Finally, embryo sac morphology was examined in two of these mutant alleles (Supplementary Figure 5 and Supplementary Table 3). In a small sample size, both alleles tested show some of the embryo sac phenotypes seen in *ig2-O/+* heterozygotes, namely the abnormal antipodal cell and extra polar nuclei phenotypes.

Taken together these data support the identity of the *ig2* gene as Zm00001d011615 (GRMZM2G030284 in B73 v3). The *ig2* gene is expressed at a much higher level in the embryo sac (16 FPKM) than the pollen grain (0.1 FPKM). The closest homolog in maize of *ig2*, Zm00001d043831 (GRMZM2G106028 in B73 v3), is in the syntenic region of chromosome 3 and has a similar expression pattern as *ig2* but at a lower level in the embryo sac (2.9 FPKM) and also very low in mature pollen (0.03 FPKM) (Chettoor et al., 2014). This is a reversal of common expression patterns as *ig2* is part of the maize subgenome 2, and Zm00001d043831 is in subgenome 1, which typically has higher expression than subgenome 2 genes (Schnable et al., 2011). The very low expression in wild-type pollen may explain why *ig2* loss-of-function affects the female gametophyte but not the male.

## DISCUSSION

Here we have described two mutants, *ig2* and *ig3*, that increase the number of polar nuclei in the central cell and cause a reduction in the number of antipodal cells. These antipodal cells are typically abnormal in morphology with them commonly being more vacuolated than in wild-type. These mutants do not appear to have excess nuclei in the syncytial phase but rather incorporate extra nuclei from the 8-nucleate syncytium into the central cell. The earliest defect seen in *ig2* is at the transition from the 8-nucleate syncytial phase to the early cellularized phase. *IG2* encodes a MAP65-3 protein which is typically localized to the phragmoplast during cytokinesis. In maize embryo sac development, the phragmoplasts are not present during the first two rounds of free-nuclear divisions and first appear after the third division just before cellularization (Huang and Sheridan, 1994). Other mutants affecting the phragmoplast, such as *atnack1* and *atnack2*, and *two-in-one*, cause embryo sac defects demonstrating the role for the phragmoplast in embryo sac cellularization (Tanaka et al., 2004; Oh et al., 2005; Takahashi et al., 2010; Sasabe et al., 2015). In Arabidopsis, there are no reported effects of mutations in the phragmoplast associated protein *MAP65-3/PLEIADE* on embryo sac cellularization (Muller et al., 2002; Ho et al., 2011; Sasabe et al., 2011), but here we show that, in maize, mutations in the *ig2/map65-3* gene cause abnormal cellularization of the female gametophyte. In the *ig2* mutant embryo sacs, not only does cellularization fail to occur normally, but the nuclei also fail to maintain their position within the syncytium. This is despite the fact that the nuclei migrate properly and maintain their position normally during the earlier phases of female gametophyte syncytial development.

Nuclear position is correlated with cell fate in the syncytial female gametophyte. This has been inferred from the phenotype of *ig1* mutants in maize (Lin, 1978, 1981; Guo et al., 2004) and several mutants in Arabidopsis (Gross-Hardt et al., 2007; Pagnussat et al., 2007; Moll et al., 2008; Kirioukhova et al., 2011; Kong et al., 2015). Manipulation of micropylar nuclear position has also been shown to alter cell fate (Sun et al., 2021). In the *ig2* mutant, the nuclei are in their normal position until the time of cellularization. Despite this, the ectopic nuclei incorporated into the central cell by the abnormal cellularization events are still able to adopt normal central cell identity. These extra polar nuclei must obtain normal central cell expression patterns for the central cell to produce viable endosperms when fertilized by the diploid sperm of tetraploid males (Figure 1E). Parental conflict theory predicts that changes in the maternal-paternal genome ratio in the endosperm leads to endosperm abortion (Haig and Westoby, 1989). Only, an endosperm with a ratio of 2 maternal genomes : 1 paternal genome from the fertilization of a tetraploid central cell by a diploid pollen grain will develop normally; the ability to get normal kernels from pollination of *ig2/+* heterozygotes by tetraploids thus demonstrates the functionality of these nuclei as polar nuclei. If these extra nuclei failed to adopt a central cell gene expression program or undergo normal imprinting characteristic of the central cell, these extra nuclei would lead to a failure either in fertilization of the central cell or an abortion in endosperm development after pollination by tetraploid males. The ability of these nuclei to change their fate so late before cellularization demonstrates either that the establishment of the identity of these nuclei occurs late, or it is not stable.

The antipodal cells in *ig2* and *ig3* mutants lose at least some of their normal traits. They are often fewer in number than in wild type and these antipodal cells can be much larger and more vacuolated than wild type. These antipodal cell clusters with larger more vacuolated cells – which are easier to identify in the live cell imaging – typically do not express fluorescent markers typical of antipodal cells, *pDR5::RFP, pPIN1::PIN1-YFP*, and *pHISTONE H1B::H1B-YFP* (although *pHIS1 H1B::H1B-YFP* was not tested in *ig3*). The lack of *DR5* and *PIN1* expression may be associated with the lack of proliferation of the antipodal cells in these mutants; this is a phenotype seen in the *Laxmidrib1* mutant of maize which also has antipodal cells that fail to proliferate properly, although they are normal in morphology (Chettoor and Evans, 2015). The *ig2* mutants, at least, also lack expression of the *HISTONE H1B* marker.Taken together these data show that the antipodal cells lack the traits of normal antipodal cells, and it is even possible that they lack antipodal cell identity. It is not yet known if they express markers typical of other cell types such as the central cell or egg cell. One thing we have not seen in these mutants is the ectopic expression of antipodal cell markers in any of the other cells of the embryo sac. So, the inclusion of nuclei from the most chalazal positions of syncytial gametophyte into the central cell does not confer antipodal cell gene expression patterns in these mutant embryo sacs.

Despite the fact that *ig3*, like *ig2*, produces embryo sacs with extra polar nuclei, a smaller antipodal cell cluster, and abnormal antipodal cells, there are some important distinctions between them. The phenotype seen in *ig2* embryo sacs of a single large cell with centrally placed nuclei (like *pleiade* root cells in Arabidopsis) is not seen in *ig3*. This may suggest that *ig3* does not have the same function in the phragmoplast as *ig2*. Additionally, *ig3* mutants also occasionally have polar nuclei located against the side wall of the central cell. This phenotype is not seen in *ig2* but is seen in a number of maternal effect mutants in maize *baseless1, baseless2, no legacy1, heirless1, superbase1*, and *maternally reduced endosperm1* (Gutierrez-Marcos et al., 2006; Chettoor et al., 2016). These mutants produce kernels with a small endosperm that is typically abnormal in morphology and often has a loose pericarp. They also commonly cause abnormal embryo development, a phenotype seen in *ig3* but not *ig2*. It is possible then that this embryo sac phenotype in *ig3* but not *ig2* is related to the embryo defect seen in *ig3* but not *ig2*.One possible explanation of the *ig3* mutant phenotype is that it is involved in nuclear movement in the embryo sac rather than cytokinesis *per se*.

## MATERIALS AND METHODS

### Plant material and growth conditions

The *ig2-O* and *ig3-O* mutations arose in maize (*Zea mays*) plants from mixed genetic backgrounds with active *Mutator* transposable elements, from either open pollinated populations (*ig2-O*) or self-pollinated populations (*ig3-O*). The ears carrying these mutations were identified based on the combination of several phenotypes: partial sterility, the production of a relatively small number (<10%) of miniature kernels, and the production of aborted kernels. The mutations were typically propagated as heterozygotes by transmission through the female and selection for miniature kernels. Mutants were back-crossed at least five generations to a variety of inbred lines before characterization of phenotypes in those backgrounds. Mutants and wild-type controls were grown side by side for each experiment either in summer field conditions or in greenhouses under long-day conditions (16 hr light:8 hr dark cycles).

### Mapping with simple sequence repeat and insertion/deletion markers

Initial map position was determined using 30 Simple Sequence Repeat markers scattered across the genome, according to the methods (Phillips and Evans, 2011). Several mapping populations were used for *ig2-O*: *ig2-O/+* B73/W23 hybrids pollinated by B73 or *ig2-O/+* B73/Mo17 hybrids pollinated by B73 or s *ig2-O/+* B73/Mo17 hybrids self-pollinated. Linkage was initially determined using the miniature kernels, *ig2-O/+*, and linkage was only detected for markers on chromosome 8. Since each cross only produces a relatively small number of miniature kernels, larger populations to further refine the *ig2* genomic interval used normal kernels (which have an unknown *ig2* genotype); from this population recombinants between the closely flanking markers *umc1858*, and *umc1960* were chosen, self-pollinated, and seed phenotypes were scored to distinguish *ig2-O/+* and *+/+* individuals. For *ig3* mapping, miniature kernels from a cross of *ig3-O/+* W23/A158 hybrid females pollinated by A158 males were used for the mapping population, and linkage was determined as for *ig2* above with linkage only seen for markers on chromosome 2. DNA was extracted from seedlings by minor modification of the method of (Saghai-Maroof et al., 1984), and PCR reactions were performed as described (Evans and Kermicle, 2001).

### Confocal microscopy of embryo sacs

Embryo sacs were analyzed from mutant heterozygous. Samples were processed in two different ways to analyze embryo sac morphology of fixed mature and immature embryo sacs either fixed in FAA without additional staining according to according to the methods of (Phillips and Evans, 2011) and visualized on a Leica SP8 (Wetzlar, Germany) laser scanning confocal microscope or double-stained with Acriflavine as a Schiff reagent and with Propidium Iodide, according to (Chettoor and Evans, 2015). Excitation was performed at 405 nm, 488 nm, and 561 nm and emission was collected at 410-480 nm, 495-555 nm, and 565-730 nm for the merged images for the unstained images, and for the double-stained samples excitations of 436 nm and 536 nm and emissions of 540 ± 20 nm and 640 ± 20 nm were used. For live-cell imaging of fluorescent reporters, *ig2-O/+* and *ig3-O/+* heterozygotes were crossed as females by reporter line hemizygotes, *pHISTONE H1B(GRMZM2G164020)::HISTONE H1B-YFP/-, pDR5::RFP/-*, or *pPIN1(GRMZM2G098643*)*::PIN1-YFP/-*. Embryo sacs were visualized from plants that were heterozygous for the mutant alleles and hemizygous for the transgenic fluorescent reporter. All transgenic lines were generously supplied by the Maize Cell Genomics Project (http://maize.jcvi.org/cellgenomics/index.php) (Mohanty et al., 2009). Miniature kernels were planted, tested for the transgene based on Basta resistance, and unpollinated ears were collected, ovules were dissected from fresh ears, and embryo sacs imaged according to (Chettoor and Evans, 2017). Images were analyzed and processed using Fiji (NIH).

### Generation of additional *ig2* mutant alleles

To produce the CRISPR-Cas9 induced ig2 mutants, guide RNA targeting the genomic site (5′-GTTGCGGCAGGCCATTGCGG<u>AGG</u>-3′ with underlined AGG as PAM) was constructed and combined with Cas9 expression destination plasmid. Cas9/gRNA was introduced into *Agrobacterium* which was used for Hi-II transformation as previously described (Char et al., 2017). Eight independent transgenic events (multiple plantlets per event) in a total of 46 plantlets were generated and screened for edits using oligonucleotides ig2-F (5′-TTTCCTGGGTCGTGTCAGG-3′) and ig2-R (5′-AATGCATGCAGTTCCTGACC-3′) to PCR-amplify the target region and sequencing of amplicons. Edited plants were grown to maturity. From these eight, we successfully recovered three heritable, mutant alleles.

## Author Contributions and Acknowledgments

Matthew Evans designed the study, identified the *ig2* and *ig3* mutant isolates, performed most of the transmission experiments, performed some of the *ig2* histology, and performed the initial mapping experiments. Antony Chettoor performed the fine-scale mapping and over 90% of the histology of *ig2* and *ig3*, both of fixed and live material. Bing Yang performed the CRISPR-Cas9 mutagenesis of the *ig2* gene to generate additional mutant alleles. The authors would like to thank Kathy Barton for the support, encouragement, and insight that kept this project alive, as well as help with the screen that identified the *ig2-O* mutant. We thank members of the Evans and Barton labs past and present for helpful discussions and thank Lance Cabalona, Clayton Coker, Amber Glowacki, Hannah Vahldick, and Michelle Pazmiño Cajiao for help growing plants and making crosses. We thank the Maize Cell Genomics project for the fluorescent reporter lines. We thank Alice Barkan and Rosalind Williams-Carrier for the *Mu* Illumina sequencing of the *ig2* and *ig3* mutant lines. This project was supported by National Science Foundation Plant Genome Program Grant, DBI-1340050.

## Supplementary Tables

**Supplementary Table 1.** Early stage embryo sac phenotypes in *ig2-O*.

| Supplementary Table 1. Early stage embryo sac phenotypes in *ig2-O* |  |  |  |  |  |
| --- | --- | --- | --- | --- | --- |
|  | 2-nucleate stage |  |  |  |  |
|  | One nucleus at each pole |  | Either the micropylar or chalazal pole lacks a nucleus |  |  |
| +/+ | 17 |  | 0 |  |  |
| *ig2-O*/+ | 33 |  | 0 |  |  |
|  | 4-nucleate stage |  |  |  |  |
|  | Chalazal nuclei pair is perpendicular to long axis of the embryo sac |  | Chalazal nuclei pair is parallel to the long axis of the embryo sac and the nuclei are adjacent |  | Chalazal nuclei pair is parallel to the long axis of the embryo sac and there is a gap between them |
| +/+ | 9 |  | 6 |  | 5 |
| *ig2-O*/+ | 21 |  | 11 |  | 7 |
|  | 8-nucleate stage/early cellularized |  |  |  |  |
|  | Not cellularized | Cellularized | Two Polar nuclei are adjacent and in the central region of embryo sac | Two polar nuclei are not adjacent | More than two nuclei in the center of the embryo sac (3-6 nuclei) |
| +/+ | 8 | 60 | 64 | 4 | 0 |
| *ig2-O*/+ | 23 | 25 | 37 | 5 | 6 |

**Supplementary Table 2.** Seed phenotypes of CRISPR alleles of *map65-3* and non-complementation of *ig2-O* by CRISPR *map65-3* alleles.

|  |  |  |  |  |  |  |  |  |
| --- | --- | --- | --- | --- | --- | --- | --- | --- |
|  | Normal endosperm |  |  | Miniature endosperm |  |  | Aborted endosperm/<br>empty pericarp | Etched, or pitted endosperm |
| Inbred background and type of cross | Normal embryo | Abnormal or aborted embryo | Twins | Normal embryo | Abnormal or absent embryo | Twins |  |  |
| *ig2/+* heterozygous female by wild type male |  |  |  |  |  |  |  |  |
| *+/ig2-2* (*insertion A*); x *+/+* (384) | 93.7% | 0.2% | 0 | 3.7% | 0 | 0 | 2.4% | 0% |
| *+/ig2-3* (*insertion T*); x *+/+* (802) | 90.1% | 0.2% | 0 | 5.3% | 0 | 0 | 4.1% | 0.2% |
| *+/ig2-4* (*insertion G*) x *+/+* (531) | 90.1% | 0.2% | 0 | 5.2% | 0 | 0 | 7.2% | 0.5% |
| *ig2-O* non-complementation |  |  |  |  |  |  |  |  |
| *+/ig2-O* x *+/ig2-2* (*insertion A*) (250) | 72.5% | 0 | 0 | 2.6% | 0 | 0 | 24.9% | 0 |
| *+/ig2-O* x *+/ig2-3* (*insertion T*) (532) | 78.4% | 0 | 0 | 1.8% | 0 | 0 | 19.8% | 0 |
| *+/ig2-O* x *+/ig2-4* (*insertion G*) (469) | 77.3% | 0 | 0 | 3.3% | 0 | 0 | 19.4% | 0 |

**Supplementary Table 3.** Summary of Mature Embryo Sac Phenotypes caused by *ig2* CRISPR alleles.

|  |  |  |  |  |  |  |  |  |  |  |  |  |  |  |  |  |
| --- | --- | --- | --- | --- | --- | --- | --- | --- | --- | --- | --- | --- | --- | --- | --- | --- |
| Normal | Abnormal |  |  |  |  |  |  |  |  |  |  |  |  |  |  |  |
|  | Aborted/<br>arrested | 1<br>Big<br>Cell | Antipodal Cell Phenotypes |  |  |  |  | Egg Apparatus Phenotypes |  |  |  |  | Central Cell Phenotypes |  |  |  |
|  |  |  | Fewer AC |  |  |  | wt<br>AC | Fewer Egg + Syn |  |  |  | Normal<br>Syn +<br>Egg | Normal<br>CC | Extra<br>PN | Extra<br>CC | 1<br>PN |
|  |  |  | Fewer<br>but<br>normal | No<br>AC | Small but<br>vacuolated | Large |  | 2<br>Syn<br>0<br>Egg | 0<br>Syn<br>1<br>Egg | 1<br>Syn<br>1<br>Egg | 1<br>Syn<br>0<br>Egg |  |  |  |  |  |
| +/ig2-3 (*insertion T*) |  |  |  |  |  |  |  |  |  |  |  |  |  |  |  |  |
| 11 | 0 | 0 | 0 | 1 | 3 | 8 | 0 | 0 | 1 | 2 | 1 | 8 | 6 | 5 | 0 | 1 |
| +/ig2-4 (*insertion G*) |  |  |  |  |  |  |  |  |  |  |  |  |  |  |  |  |
| 8 | 0 | 0 | 0 | 0 | 0 | 6 | 0 | 1 | 0 | 0 | 1 | 4 | 0 | 1 | 0 | 1 |

**Supplementary Figure 1.**
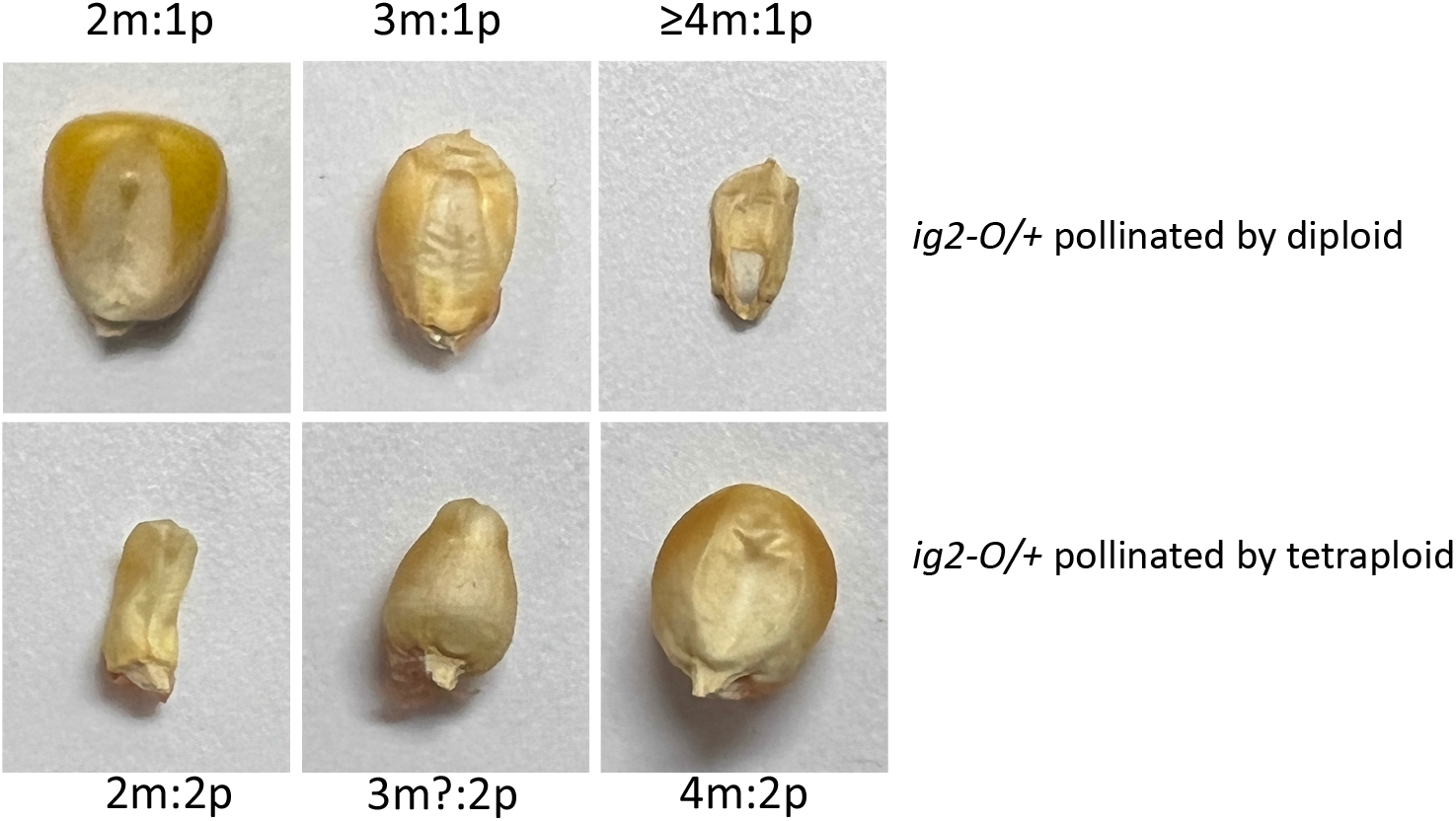
Kernels produced by *ig2-O/+* heterozygotes when crossed by diploid or tetraploid males. A range of kernel sizes are produced when *ig2* or *ig3* mutants are crossed by diploid or tetraploid males. The number of maternal genomes in the endosperm varies with the number of polar nuclei in the central cell. The maternal genome dosage here is inferred from the kernel phenotype.

**Supplementary Figure 2.**
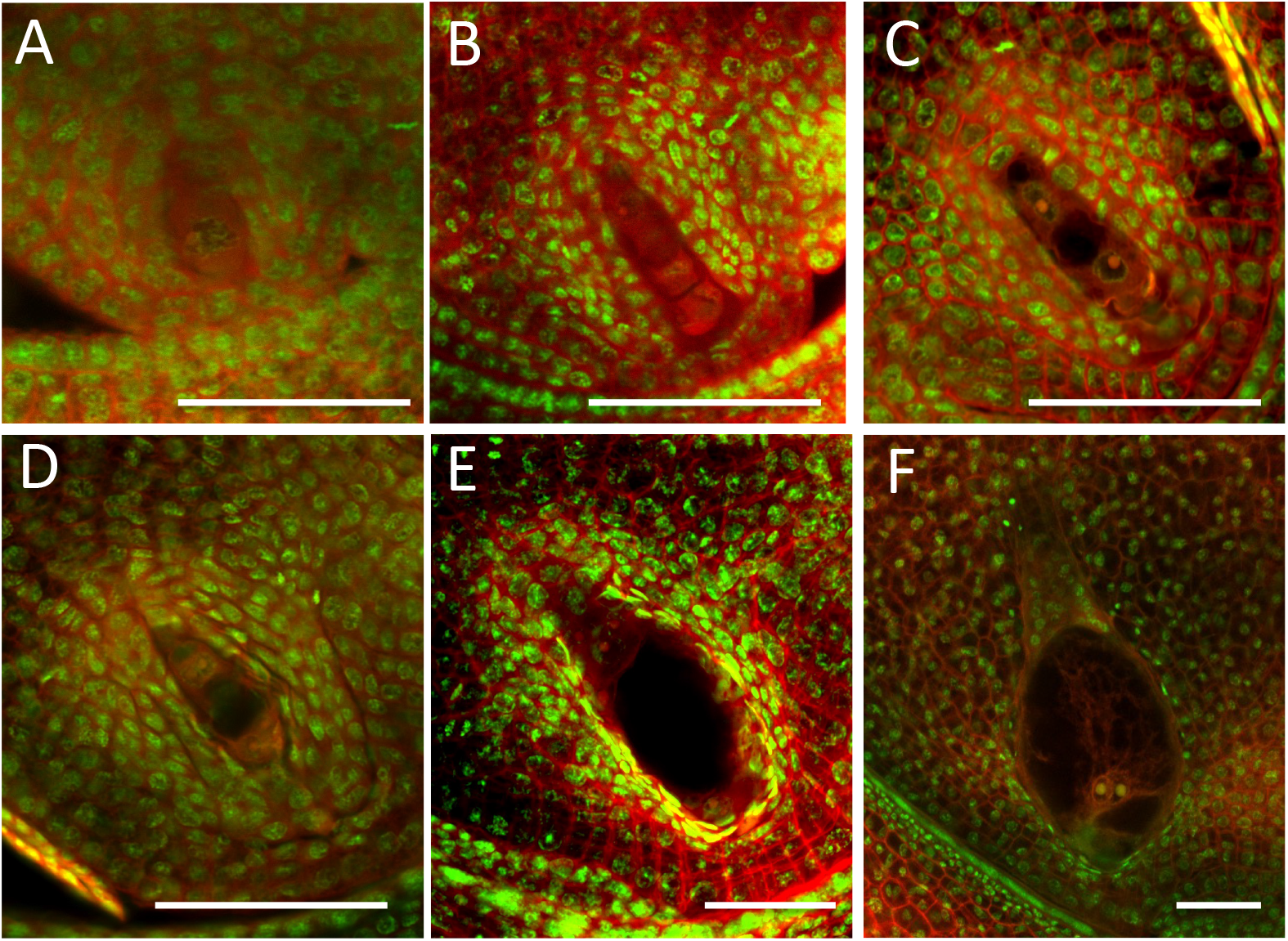
Wild type embryo sac developmental time course. (A) Megaspore Mother Cell. The nucleus has already changed in morphology from the surrounding nucellus with amore prominent nucleolus and reduced Propidium Iodide staining. (B) Functional megaspore/one-nucleate female gametophyte with degenerating micropylar megaspores. (C) Two-nucleate female gametophyte. (D) Four-nucleate female gametophyte. (E) Eight-nucleate female gametophyte. (F) Mature female gametophyte Scale bar=50μm

**Supplementary Figure 3.**
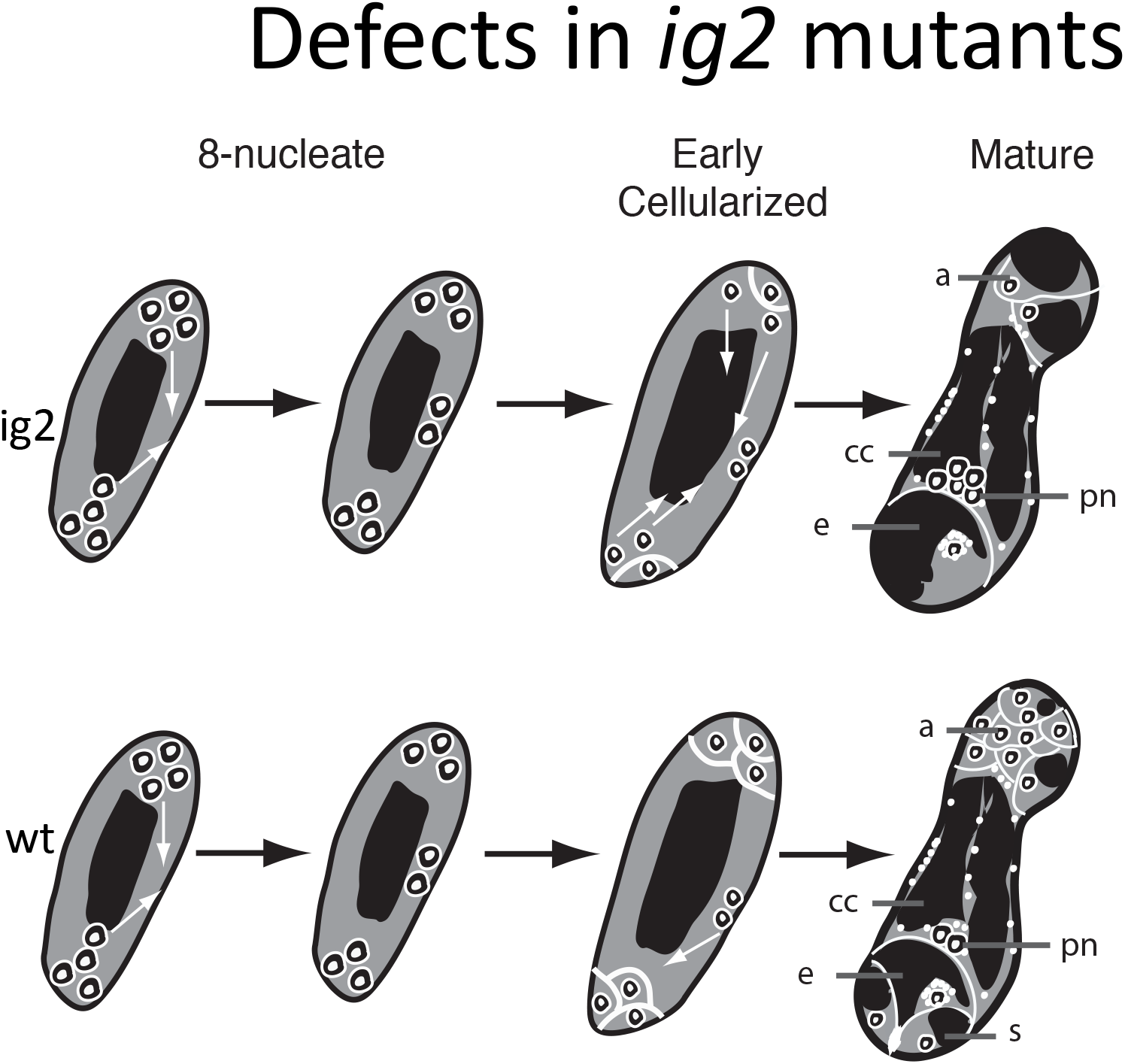
Schematic of developmental defects in *ig2* mutant embryo sacs.

**Supplementary Figure 4.**
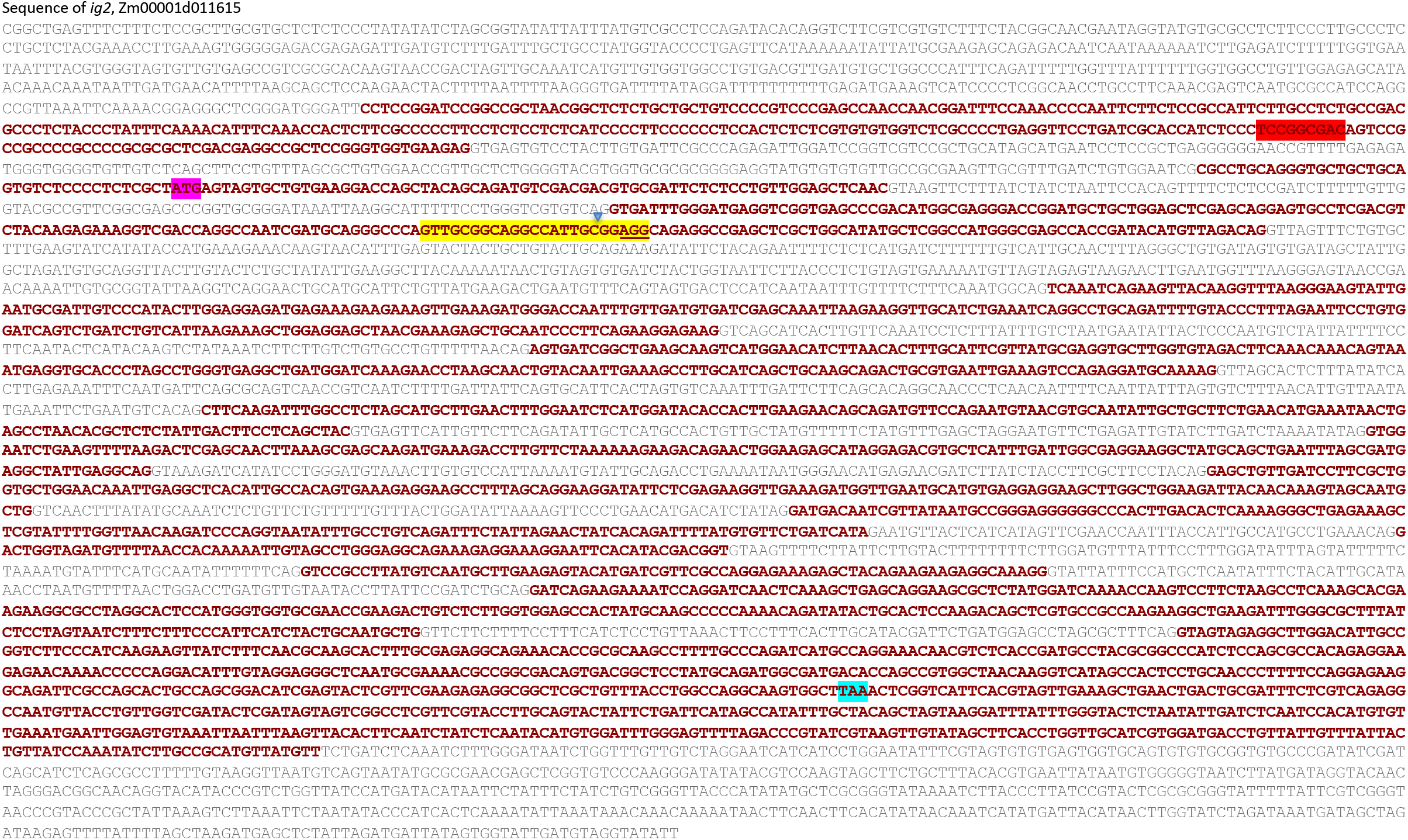
*ig2* gene structure. Exons are marked with maroon letters, and the start and stop codons are highlighted in magenta and cyan, respectively. The target site duplicated in the *Mu* insertion of *ig2-O* is highlighted in red. The CRISPR target sequence is highlighted in yellow. The position of the single nucleotide insertion alleles produced by the CRISPR/CAS9 targeting is marked with a blue triangle. Three different alleles were recovered from the CRISPR targetting: one with an insertion of an A, one with a G, and one with a T, all at the same position.

**Supplementary figure 5.**
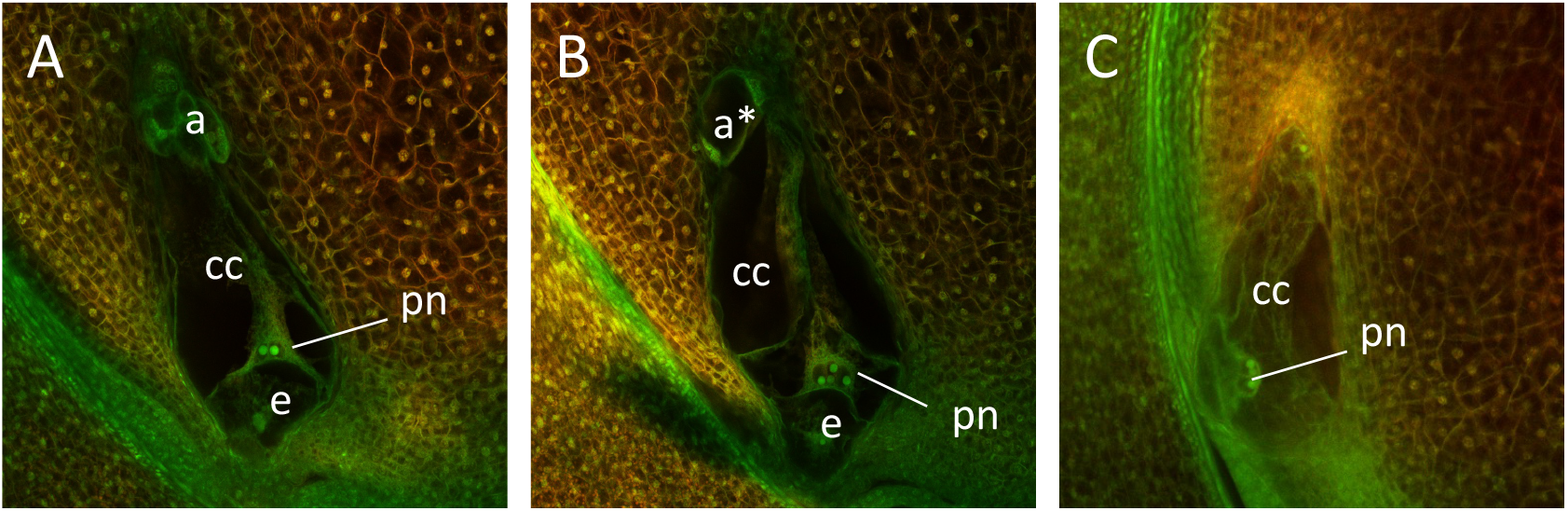
Embryo sac phenotypes from heterozygotes for *ig2-4* (G insertion), one of the CRISPR-induced mutant alleles of *ig2*. (A) Normal embryo sac. (B,C) *ig2-4* abnormal. (B) Abnormal embryo sac with enlarged vacuolated antipodal cells and extra polar nuclei. (C) Abnormal embryo sac with no antipodal cells.

